# ERK signaling integrates multiple mechanisms to drive cortical evolutionary expansion

**DOI:** 10.64898/2026.07.28.741115

**Authors:** Tongye Fu, Zhuangzhi Zhang, Wenhui Zheng, Chuannan Yang, Zizhuo Sha, Danyu Han, Jialin Li, Feihong Yang, Jingzhe Yu, Zhenmeiyu Li, Yan You, Tong Ma, Weiwei Li, Guoping Liu, Xiaolei Song, Dashi Qi, Da Mi, Wei Huang, Zhejun Xu, Zhengang Yang

## Abstract

A critical first step in human evolutionary divergence from chimpanzees and other primates is the production of greatly increased numbers of cortical pyramidal neurons during development and over evolutionary time. This expansion underpins the enlarged human cerebral cortex and, consequently, higher-order cognition and consciousness. However, the cellular and molecular mechanisms driving this progressive increase in cortical neuron number remain incompletely understood. Here we show that elevated ERK signaling in human cortical radial glia (RGs), relative to mouse RGs, arises from evolutionary changes in the developmental expression of existing shared genes. Using the *Emx1-Cre* line to overexpress *MEK1DD*, a constitutively active mutant of rat *MAP2K1*, we found that enhanced ERK signaling in mouse cortical RGs upregulates cAMP–PKA signaling, along with promoting self-renewal, expanding the RG pool, and accelerating cell cycle progression. Conversely, overexpression of PKA signaling in mouse cortical RGs via in utero electroporation of constitutively active *PRKACA* mutants (L206R or W196G) reduces ERK activity, prolongs the cell cycle, suppresses SHH–SMO and YAP/TAZ pathway activity, and blocks ependymal gliogenesis. Together, our results demonstrate that ERK and PKA signaling in cortical RGs engage a dynamic balance of synergistic and antagonistic interactions. This interplay couples evolutionarily enhanced ERK and PKA activities to increased cortical RG numbers, prolonged cell cycle progression, and an extended neurogenic period, which collectively culminate in a marked increase in neuronal production. We conclude that heightened ERK pathway activity in cortical RGs serves as a central driver of progressive neocortical expansion across mammalian evolution.

**Significance Statement:** The human brain’s extraordinary cognitive abilities stem from a dramatic increase in cortical neuron numbers during evolution. This study reveals that enhanced ERK signaling in human cortical radial glia neural stem cells, relative to other primates, is a key driver of this expansion. By manipulating this pathway in mice, we demonstrate that ERK coordinates multiple mechanisms—including self-renewal, cell cycle acceleration, and a dynamic interplay with PKA signaling—to boost neuronal output. These findings identify a core molecular axis underlying mammalian cortical evolution and provide a framework for understanding how the human cortex acquired its unique size and, consequently, its remarkable complexity.

## Introduction

During early pallium development, neuroepithelial cells expand their progenitor pool through symmetric proliferative divisions. As the pallium thickens, these cells progressively transform into radial glia (RGs), which serve as the primary neural stem cells (1–12). Concurrently, cortical RG divisions shift from symmetric to asymmetric modes. This shift occurs from a ventrolateral to a dorsal position, during which cortical RGs undergo self-renewal while generating one intermediate progenitor cell (IPC) per division. Thus, most cortical pyramidal neurons (PyNs) and glial cells are not direct progeny of RGs, but rather arise from these transit-amplifying IPCs (1–12). In mice, cortical RGs retain their characteristic full-span morphology throughout cortical development. In humans, full-span radial glia (fRGs) persist from gestational week (GW) 8 to GW16, similar to their murine counterparts. Thereafter, human cortical fRGs gradually bifurcate, producing truncated radial glia (tRGs) in the ventricular zone (VZ) and outer radial glia (oRGs, also known as basal radial glia) in the outer subventricular zone (OSVZ) (13–20).

Mammalian cortical RGs produce their daughter IPCs via an intrinsically programmed temporal cascade. This cascade initially yields exclusively PyN-IPCs and later shifts toward gliogenesis—a process that produces cortical ependymal cells, astrocytes, and oligodendrocytes. Consequently, at any given stage of cortical development, every RG exhibits a distinct intrinsic bias toward generating a specific class of progeny. In the mouse cortex, RGs initially undergo neurogenesis and are termed neurogenic RGs (N-RGs) (21, 22). Following cortical neurogenesis, a subset of N-RGs in the medial cortex are the first to convert into E-RGs, which almost exclusively generate ependymal cells (21–26). In contrast, the other subset of N-RGs progressively transitions into T-RGs along a ventrolateral-to-dorsolateral cortical gradient. T-RGs generate a distinct type of IPC, now increasingly recognized as cortical tripotential IPCs (Tri-IPCs), which in turn sequentially produce three classes of fate-restricted IPCs: APCs for astrocytes, OPCs for oligodendrocytes, and OBIN-IPCs for cortically derived olfactory bulb interneurons (4, 12, 16, 21, 22, 27–42). It is important to note that the term Tri-IPC does not imply that each Tri-IPC must generate APCs, OPCs, and OBIN-IPCs *in vivo* during normal cortical development (43). Rather, an individual Tri-IPC may produce only one or two fate-restricted IPC types, yet the term denotes that Tri-IPCs as a population genuinely possess tripotent differentiation capacity. This point is clearly illustrated by loss-of-function experiments. Notch inhibition restricts Tri-IPC output to OPCs and OBIN-IPCs while abolishing APC production (29, 35). Conversely, loss of *Olig1/2* shifts differentiation toward APCs and OBIN-IPCs at the expense of OPCs (32, 34). In contrast, *Gsx1/2* or *Dlx1/2* deficiency permits APC and OPC formation but eliminates OBIN-IPCs (44–46). The tripotent potential of single neural stem cells and IPCs derived from the adult mouse SVZ has been also demonstrated by in vitro clonal analysis using neurosphere assays or adherent cultures, as well as by *in vivo* intraventricular EGF injection (28, 47, 48).

In the human cortex, fRGs are neurogenic and generate deep-layer PyNs. After tRGs and oRGs are established around GW16, oRGs continue to sustain the neurogenic function of fRGs and predominantly generate upper-layer PyNs (4, 16, 19, 21, 22, 29, 31, 49). Human cortical tRGs undergo a developmental switch to acquire gliogenic potential. Specifically, tRG subsets in the medial cortex first adopt an E-tRG identity to produce ependymal cells (16, 21, 22, 28, 50–52), whereas their ventrolateral-dorsal counterparts first transition into a T-tRG fate, giving rise to Tri-IPCs (4, 16, 19, 21, 22, 28, 29, 31, 36).

Radial glia (RGs) were first observed in the late 19th century (53, 54). In the early 1970s, Pasko Rakic introduced the term “radial glia” and established their role as a transient scaffolding population for neuronal migration (55, 56). This view was revised in the early 2000s, when multiple laboratories demonstrated that rodent RGs generate neurons and function as primary neural stem cells (57–61). Building upon the above-mentioned identification of distinct cortical RG subtypes in gyrencephalic species such as ferrets, macaques, and humans (13–17, 20–22, 49, 50, 62), a key question is to clarify their interrelationships and molecular characteristics. Indeed, our recent cross-species studies in mice, ferrets, and humans have refined the molecular profiles of three cortical RG populations: (1) PyN-generating N-RGs (mouse N-RGs; human fRGs and oRGs); (2) ependymal-generating E-RGs (mouse E-RGs; human E-tRGs); and (3) Tri-IPC-generating T-RGs (mouse T-RGs; human T-tRGs) (21, 22). Specifically, we found that the relative activities of three signaling pathways determine the subtype identity of cortical RGs and thereby dictate the fates of their progeny. N-RGs engage ERK/PKA signaling, which concurrently represses both YAP/TAZ and SHH–SMO activities. By contrast, E-RGs depend on YAP/TAZ signaling, which suppresses PKA, ERK, and SHH–SMO. T-RGs exhibit progressively increasing SHH–SMO signaling, which represses ciliary PKA while exerting only weak repression on ERK. These three pathways—ERK/PKA, YAP/TAZ, and SHH– SMO—thus constitute a tripartite regulatory network in which cross-repressive interactions coordinate distinct aspects of neurogenesis and gliogenesis (21, 22). Here, we formally define this discovery as the cortical radial glia tri-lineage principle.

The common ancestor of humans and mice lived approximately 90 million years ago. By that time, the basic blueprint of the mammalian brain—characterized by a six-layered neocortex—had already been established. Our recent studies suggest that elevated ERK activity in cortical RGs serves as a key driver of mammalian cortical expansion (19, 21, 22, 29, 31). To explore the mechanisms underlying elevated ERK activity in human cortical RGs, we performed cross-species comparative analyses.

Heightened ERK signaling in human versus mouse RGs arises from evolutionary changes in conserved gene expression. *Emx1-Cre*–mediated expression of constitutively active *MEK1DD* in mouse RGs enhanced ERK signaling, promoting RG self-renewal and expansion, accelerating cell cycle progression, and upregulating cAMP–PKA. Conversely, forced high-intensity PKA activation attenuated ERK, delayed cell cycle, repressed SHH–SMO and YAP/TAZ, and blocked Tri-IPC and ependymal cell generation. Thus, ERK and PKA exhibit both synergistic and antagonistic interactions, linking enhanced ERK/PKA activities to expanded RG pools, slower cell cycles, prolonged neurogenesis, and robust cortical PyN production.

## Results

### Human cortical fRGs express many ERK-boosting genes that are low or absent in mouse

Our recent studies demonstrate markedly higher ERK signaling pathway activity in human cortical RGs than in their mouse counterparts (19, 21, 22, 29, 31). To further investigate why human cortical fRGs exhibit higher ERK activity than mouse RGs at comparable developmental stages (human GW12 versus mouse E12.0) (63), we performed a cross-species analysis of the scRNA-seq data using a common orthologous gene space (**Fig. 1*A***). This revealed approximately 8,000 differentially expressed genes (DEGs) between cortical RGs of the two species, of which 4,700 were downregulated and 3,300 upregulated in human. We next performed KEGG pathway analysis on DEGs in human GW12 versus mouse E12.0 cortical RGs. This revealed that ERK signaling (MAPK and Ras) and cAMP-PKA signaling (cAMP, Rap1, and thyroid hormone) pathways were significantly enriched in human GW12 full-span (fRGs), whereas cell cycle, ribosome, and Hippo signaling genes were more highly expressed in mouse RGs (**Fig. 1*B***).

**Fig. 1.**
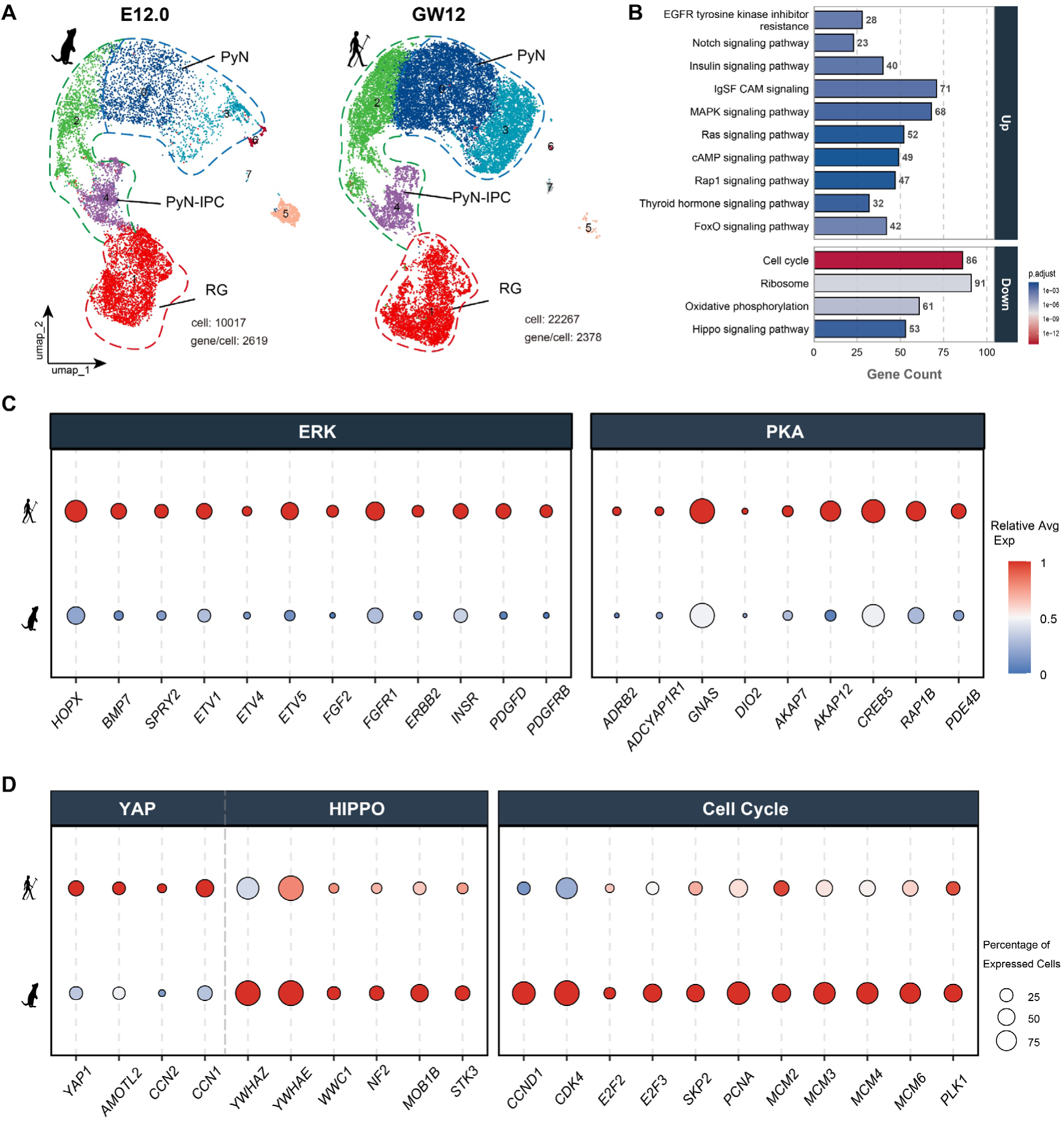
Human cortical full-span RGs exhibit elevated ERK and cAMP-PKA signaling relative to mouse RGs at equivalent stages. (*A*) UMAP projection of cross-species scRNA-Seq data from human GW12 and mouse E12.0 cortices, colored by cluster identity. PyN, cortical glutamatergic pyramidal neurons; PyN-IPC, PyN intermediate progenitor cells; RG, radial glia. (***B***) KEGG pathway analysis of genes significantly up- and downregulated in GW12 human cortical RGs vs. E12.0 mouse cortical RGs. (***C, D***) Bubble plot illustrating differentially expressed genes (DEGs) between human GW12 and mouse E12.0 cortical RGs. Dot size reflects the percentage of cells expressing the gene, while color intensity indicates the maximum-normalized relative expression level. Human GW12 RGs show elevated ERK, cAMP-PKA, and YAP/TAZ signaling, whereas mouse E12.0 RGs exhibit enhanced Hippo signaling (suppressing YAP/TAZ) yet faster cell cycle progression.

KEGG and GO pathway databases primarily catalog direct signaling components, such as enzymes and functional proteins, rather than downstream effectors (readout genes). To more rigorously determine whether these pathways are genuinely activated or suppressed, we examined both core components and known readout genes. Indeed, human cortical fRGs expressed higher levels of *HOPX*, *BMP7*, *SPRY2*, and *ETV1/4/5*—all well-established ERK signaling readouts in cortical RGs. Furthermore, human cortical fRGs also showed elevated expression of *PDGFD*, *PDGFRB*, *INSR*, *ERBB2*, *FGF2*, and *FGFR1*, which drive ERK signaling activation. In contrast, expression of these genes in mouse cortical RGs was low or undetectable (**Fig. 1*C*** and *SI Appendix,* Fig. S1). These species-specific expression patterns suggest that evolutionary innovation in cortical RG development arises not from new genes, but from rewiring of conserved regulatory networks—a mechanism that drives the markedly higher ERK signaling in human RGs and carries significant evolutionary implications. Similarly, this mechanism also applies to cAMP-PKA and YAP signaling, both of which are much more active in human cortical RGs, whereas mouse cortical RGs show higher Hippo signaling (which represses YAP/TAZ) and increased cell cycle gene expression (**Fig. 1*C*, *D*** and *SI Appendix,* Fig. S2).

Notably, although PKA and YAP/TAZ signaling repress each other in cortical RGs, human cortical fRGs exhibit higher levels of both pathways than mouse RGs. This apparent paradox is resolved by regional heterogeneity: ventral–lateral–dorsal cortical RGs show elevated ERK signaling, whereas medial cortical RGs show elevated YAP compared with their mouse counterparts (21, 22). These findings suggest that gliogenic signals—including YAP and SHH–SMO—must be progressively upregulated to overcome the predominance of neurogenic signals in cortical N-RGs, thereby enabling the initiation of cortical gliogenesis at later developmental stages. Otherwise, persistently high ERK/PKA signaling alone would preclude the elevation of YAP/TAZ and consequently impair ependymal glial cell generation.

Taken together, our results demonstrate that the heightened ERK, cAMP-PKA, and YAP/TAZ signaling—all evolutionarily ancient pathways—in human cortical RGs is predominantly attributable to network-level rewiring rather than the deployment of new genes.

### Human cortical oRGs exhibit elevated ERK/PKA signaling over tRGs from their emergence at GW16

Human cortical fRGs begin to generate—or divide into—tRGs and oRGs around GW16, coinciding with the gradual disappearance of the fRG population itself (15). We therefore performed pERK (phosphorylated ERK) immunohistochemistry on human GW16 cortex. This analysis revealed that oRGs already express visibly higher pERK levels than tRGs at their emergence (**Fig. 2*A, B***).

**Fig. 2.**
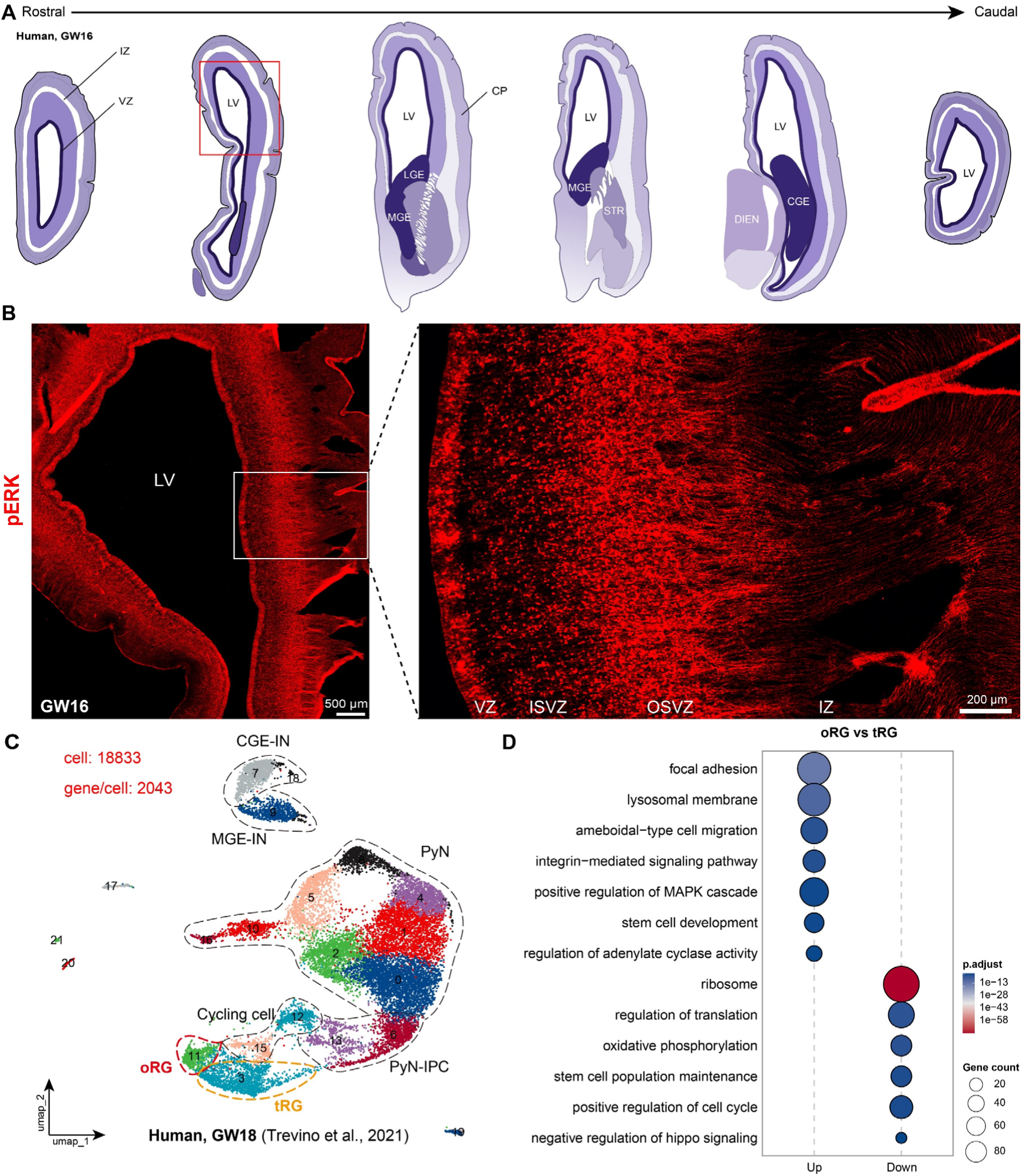
Human cortical oRGs display elevated ERK signaling compared to tRGs as early as GW16, the time of their emergence. (*A*) Human telencephalon coronal sections along the rostral–caudal axis at GW16. CGE, LGE, MGE, caudal, lateral, medial ganglionic eminence; CP, cortical plate; DIEN, diencephalon; IZ, intermediate zone; LV, lateral ventricle; STR, striatum; VZ, ventricular zone. (***B***) GW16 coronal section of the rostral telencephalon (boxed in ***A***). pERK immunoreactivity was stronger in the OSVZ than in the VZ. ISVZ, inner subventricular zone. (***C***) Re-analysis of published GW18 human cortical scRNA-Seq data (36). Cortical RGs were simply classified into oRGs and tRGs. (***D***) Gene Ontology (GO) term analysis of DEGs between oRGs and tRGs reveals enrichment of ERK signaling (positive regulation of MAPK cascade) and cAMP-PKA signaling (regulation of adenylate cyclase activity) in human cortical oRGs. Dot size represents the number of DEGs enriched in each term, and color intensity indicates the adjusted P value.

We next analyzed published scRNA-Seq data from human cortical GW18 (36), the nearest available developmental time point to GW16. At GW18, human cortical RGs were classified simply as oRGs and tRGs (**Fig. 2*C***). Our analysis revealed approximately 2,000 DEGs between cortical oRGs and tRGs, of which 1,300 were upregulated and 700 downregulated in human cortical oRGs relative to tRGs. We then performed Gene Ontology (GO) term analysis on DEGs in oRGs versus tRGs. This revealed that transcripts involved in ERK signaling (positive regulation of MAPK cascade) and cAMP-PKA signaling (regulation of adenylate cyclase activity) were enriched in human GW18 cortical oRGs, whereas cell cycle (positive regulation of cell cycle), ribosome, and YAP/TAZ signaling (negative regulation of Hippo signaling) pathway genes were more highly expressed in human cortical tRGs (**Fig. 2*D***).

Another key indicator of elevated ERK signaling in human oRGs is *HOPX* expression, which is almost entirely dependent on ERK signaling: HOPX was eliminated in mouse cortical RGs lacking *Map2k1* and *Map2k2* (*SI Appendix,* Fig. S3) (21, 22, 29, 31), the core genes of the ERK pathway. Thus, the high HOPX expression in human oRGs reflects their elevated ERK activity (*SI Appendix,* Fig. S4) (31).

We also noted that, in the GW16 human medial cortex—likely due to high YAP/TAZ activity (21, 22)—SOX9⁺ cortical RGs generate few, if any, IPCs and lack both the OSVZ and oRGs (*SI Appendix,* Fig. S5). In contrast, in the dorsal and lateral cortex at GW16, both tRGs and oRGs expressing SOX9 generate abundant IPCs (*SI Appendix,* Fig. S5). Similarly, at GW23, SOX2⁺/FOXJ1⁺ E-tRGs were identified in the medial cortex, which lacked IPCs, OSVZ, and oRGs (*SI Appendix,* Fig. S6). By contrast, in the ventral and lateral cortex, SOX2⁺ T-tRGs (FOXJ1-negative), IPCs, and oRGs were clearly observed (*SI Appendix,* Fig. S6).

Taken together, our findings demonstrate that human oRGs acquire elevated ERK/PKA signaling upon emergence. The spatial balance between ERK/PKA and YAP/TAZ is critical for proper cortical developmental timing. The early rise in ERK/PKA promotes oRG proliferation, self-renewal, and extended neurogenesis while suppressing gliogenesis, thereby supporting massive PyN production.

Conversely, high YAP/TAZ activity in the medial cortex correlates with reduced IPC and oRG abundance and drives earlier generation of the E-tRG–ependymal lineage (19, 21, 22, 29, 31).

### JUNB does not bidirectionally promote cortical neurogenesis

A recent study reported that JUNB bidirectionally promotes cortical RG proliferation and neurogenic timing in humans, but not in mice, thereby augmenting overall cortical neuron production (64). To investigate this species-specific difference, we first examined *Junb* expression in mouse cortical RGs across E15.0, E17.5, P0, and P2 using scRNA-seq datasets. *Junb* was barely detectable before E17.5—the neurogenic stage—but emerged at P0 and became strongly enriched by P2—the stage at which RGs mainly give rise to ependymal cells and OB interneurons (*SI Appendix,* Fig. S7, S8). KEGG pathway analysis comparing P2 and E15.0 RGs revealed significant enrichment of the TNF signaling pathway (mmu04668, adjusted P = 0.00045) in P2 RGs, with *Junb* as a component gene (*SI Appendix,* Fig. S8*D*). GO analysis of *Junb*-associated genes upregulated in P2 versus E15.0 RGs showed functional enrichment in transcriptional regulation, immune differentiation, and responses to hormonal and ionic stimuli (*SI Appendix,* Fig. S8*E*). *Junb* was also highly expressed in postnatal cortical Tri-IPCs and cycling IPCs, both of which also exhibited high *Egfr* levels. In *hGFAP-Cre; Egfr-cko* mice at P2, *Junb* and *Egr1* expression were both markedly reduced (*SI Appendix,* Fig. S9), indicating that *Junb* expression in postnatal cortical IPCs is largely *Egfr*-dependent.

We next examined *JUNB* expression in the human cortical RGs at GW12, GW14, GW17, GW18, GW22, GW23, and GW26. *JUNB* was barely detectable at GW12 and GW14, when cortical fRGs are engaged exclusively in neurogenesis (*SI Appendix,* Fig. S10). By GW17–GW18, human cortical RGs have diverged into oRGs and tRGs. We detected marked *JUNB* expression in tRGs, while only minimal levels were observed in neurogenic oRGs at these stages (*SI Appendix,* Fig. S11, S12). At GW22, GW23, and GW26, *JUNB* expression was lowest in oRGs, moderate in T-tRGs, and highest in E-tRGs and Tri-IPCs (*SI Appendix,* Fig. S13, S14). KEGG pathway analysis of scRNA-seq data at GW22–GW26 showed that the TNF signaling pathway was significantly enriched in E-tRGs compared with oRGs, and JUNB was among the constituent genes (*SI Appendix,* Fig. S14*C*). GO analysis of *JUNB*-associated genes upregulated in E-tRGs versus oRGs revealed functional enrichment in transcriptional regulation, immune differentiation, and responses to hormonal and ionic stimuli (*SI Appendix,* Fig. S14*D*)—a pattern remarkably similar to that observed for murine *Junb* (*SI Appendix,* Fig. S8*E*).

Taken together, our findings indicate that JUNB does not play a major role in cortical neurogenesis in either species, consistent with its predominant expression in gliogenic RGs. In contrast, we demonstrate that human cortical fRGs (GW8–GW16) and oRGs (GW16–GW26) sustain elevated ERK/PKA signaling throughout this developmental window (*SI Appendix,* Fig. S10-S14), strongly supporting the notion that ERK serves as the central driver of RG proliferation, self-renewal, and the extended neurogenic period that characterizes human cortical development (19, 21, 22, 29, 31).

### ERK overexpression in mouse early cortical RGs partially recapitulates human neurogenic RG properties

Human cortical fRGs exhibit much higher ERK signaling than their mouse counterparts. To test the functional impact of elevated ERK signaling in vivo, we overexpressed a constitutively active rat *MEK1DD* mutant in mouse cortical RGs using the *Emx1-Cre* driver from E10.0 onward. We previously performed bulk RNA-seq on E14.5 cortices from *hGFAP-Cre; Rosa26^MEK1DD/+^*mice (31). Here, we extended this analysis with scRNA-seq on E12.0 control and *Emx1-Cre; Rosa26^MEK1DD/+^* cortices, focusing specifically on RGs (**Fig. 3*A, B***). This approach provides substantially higher resolution than bulk RNA-seq and represents the first scRNA-seq analysis of this model, offering RG-specific insights into the transcriptional consequences of *MEK1DD* overexpression. Our scRNA-seq analysis revealed 2,300 DEGs between *Emx1-Cre; Rosa26^MEK1DD/+^* and control cortical RGs, of which 1,500 were significantly upregulated and 900 significantly downregulated in the mutant.

**Fig. 3.**
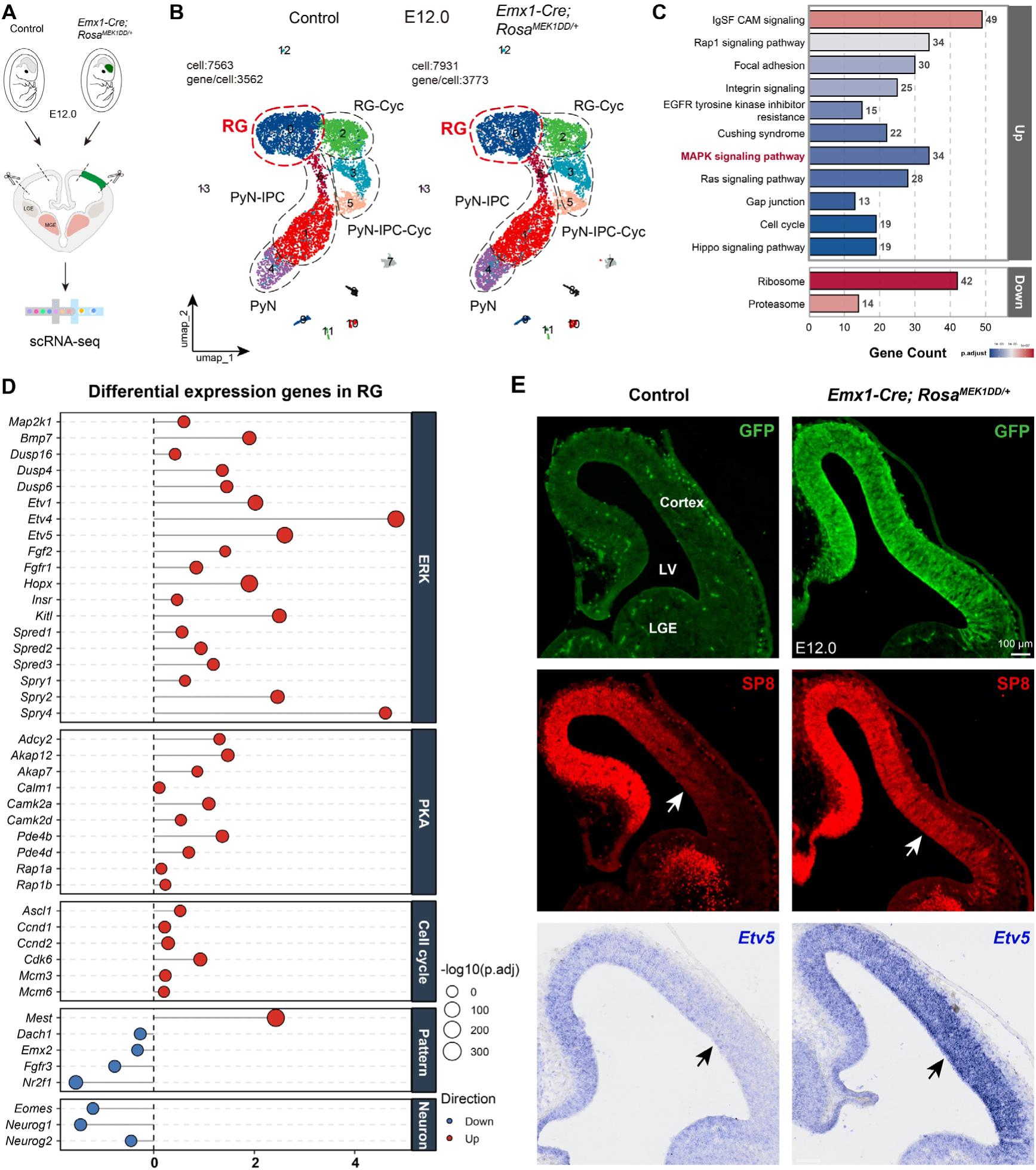
Cortical RGs in E12.0 *Emx1-Cre; Rosa^MEK1DD/+^* mice show upregulated cAMP–PKA signaling, self-renewal, RG pool expansion, and accelerated cell cycle progression. (*A, B*) Cortical scRNA-Seq analysis from E12.0 control and E12.0 *Emx1-Cre; Rosa^MEK1DD/+^* mouse embryos. (*C*) KEGG pathway analysis of genes significantly up- and downregulated in E12.0 *Emx1-Cre; Rosa^MEK1DD/+^* mouse cortical RGs compared to controls. (*D*) Faceted lollipop plot of DEGs in cortical RGs between E12.0 *Emx1-Cre; Rosa^MEK1DD/+^* and controls. Dot size indicates statistical significance (−log₁₀ adjusted *P* value). Note that *MEK1DD* overexpression in *Emx1-Cre*⁺ RGs from E10.0 upregulated *Mest* and downregulated *Dach1, Emx2, Fgfr3*, and *Nr2f1* (*Coup-TFI*), indicating rostral cortical expansion and concurrent caudal atrophy. (*E*) Immunostaining and mRNA *in situ* hybridization revealed upregulation of MEK1DD-GFP, SP8, and *Etv5*.

We next performed KEGG pathway analysis on DEGs in *Emx1-Cre; Rosa26^MEK1DD/+^* versus control cortical RGs. This revealed significant enrichment of pathways related to ERK/MAPK and Ras signaling, cAMP-PKA signaling (including Rap1 and Cushing syndrome), cell cycle, and Hippo signaling—which represses YAP/TAZ activity— among the DEGs (**Fig. 3*C***). Notably, enrichment of the Cushing syndrome pathway (mmu04934) is particularly informative, as this pathway is associated with constitutive PKA activation that drives adrenal cortisol overproduction. Its enrichment in *MEK1DD*-overexpressing RGs therefore supports the conclusion that elevated ERK signaling promotes PKA pathway activity in cortical RGs.

To validate pathway activation, we examined both core components and known readout genes. *MEK1DD* overexpression significantly upregulated *Map2k1* (encoding MEK1DD itself), *Bmp7, Dusp4/6/16, Etv1/4/5, Fgf2, Fgfr1, Hopx*, *Insr, Kitl, Spred1/2/3,* and *Spry1/2/4* (**Fig. 3*D***). This expression profile indicates that ERK overactivation in cortical RGs reshapes the signaling network by dampening negative feedback regulators (*Spry*, *Spred*, and *Dusp* families) while enhancing pro-growth signals (*Fgf2/Fgfr1/Insr*) and downstream transcriptional effectors (*Etvs, Hopx*, and *Bmp7*).

*MEK1DD* overexpression also remodels the cAMP-PKA signaling network in cortical RGs, as evidenced by upregulation of *Akap12, Akap7, Calm1, Camk2a, Camk2d*, *Pde4b, Pde4d*, and *Rap1a/Rap1b* (**Fig. 3*D***). This transcriptional reprogramming enhances PKA signaling efficiency: AKAPs organize discrete signaling microdomains, CAMs potentiate calcium signaling, and RAPs expand downstream output. Simultaneously, upregulated PDE4s activate negative feedback brakes that constrain the intensity and spatiotemporal spread of cAMP signals, thereby preventing overactivation.

At the regional level, *MEK1DD* overexpression drives frontal cortical expansion, as indicated by *Mest* upregulation in cortical RGs, while concurrently promoting caudal cortical regression and atrophy, accompanied by significant downregulation of *Dach1, Emx2, Fgfr3*, and *Nr2f1* (*Coup-TFI*) (**Fig. 3*D***). In addition, MEK1DD overexpression accelerates the cortical RG cell cycle, effectively increasing RG proliferation while preserving their self-renewal capacity. This, in turn, leads to downregulation of neurogenic genes—including *Eomes, Neurog1*, and *Neurog2*—in cortical RGs, thereby restraining neuronal differentiation (**Fig. 3*D***).

We next performed immunohistochemistry and mRNA *in situ* hybridization to confirm medial frontal cortical expansion, as evidenced by SP8 expression extending laterally into cortical RGs, and to validate *Etv5* mRNA upregulation as a readout of ERK signaling (**Fig. 3*E***). HOPX expression in control cortical RGs exhibited a rostral-high-to-caudal-low gradient, further supporting the rostrocaudal gradient of ERK signaling activity in the normally developing cortex (*SI Appendix,* Fig. S15). Notably, *Emx1-Cre; Rosa26^MEK1DD/+^* cortical RGs showed upregulated HOPX expression across the rostral-to-caudal axis from E12.0 to P5 (*SI Appendix,* Fig. S15).

We next performed a cross-species comparison of scRNA-Seq data from E12.0 control and *Emx1-Cre; Rosa26^MEK1DD/+^* mouse cortical RGs with E62/E64 macaque and GW12 human cortical RGs within a strict one-to-one orthologous gene space. Neighbor-joining and principal component analyses of these datasets suggested that *MEK1DD* overexpression partially shifts the mouse cortical RG transcriptional landscape toward that of primate cortical RGs (*SI Appendix,* Fig. S16*A-C*). SP8 immunofluorescence revealed stronger and more widespread SP8 expression in E55 macaque cortical RGs than in E12.0 control and *Emx1-Cre; Rosa26^MEK1DD/+^* mouse cortical RGs, further supporting higher ERK signaling activity in macaque cortical RGs (*SI Appendix,* Fig. S16*D*).

YAP/TAZ and ERK/PKA signaling exhibited mutual inhibition, such that YAP/TAZ-driven ependymal differentiation was accompanied by complete attenuation of ERK/PKA signaling in immature ependymal cells (*SI Appendix,* Fig. S17) (21, 22). Conversely, sustained ERK activation in *Emx1-Cre; Rosa26^MEK1DD/+^* cortical RGs completely blocked ependymal cell generation (*SI Appendix,* Fig. S18). Moreover, EGFR-ERK signaling was progressively potentiated by *MEK1DD* overexpression in *Emx1-Cre; Rosa26^MEK1DD/+^* RGs during later development, driving gliogenesis and promoting the proliferation of APC, OPC, and OBIN-IPCs (*SI Appendix,* Fig. S19) (29, 31).

Together, these findings demonstrate that ERK signaling in cortical RGs exerts broad and multifaceted effects. In early cortical development, it upregulates cAMP-PKA signaling, promotes self-renewal, expands the RG pool, and accelerates cell-cycle progression. During late cortical development, elevated ERK signaling activates PKA, which in turn upregulates Hippo signaling and suppresses YAP/TAZ activity, thereby blocking ependymal cell genesis. Sustained high ERK activity also potentiates EGFR-ERK signaling, promoting the proliferation of APCs, OPCs, and OBIN-IPCs in the late cortex.

### PKA overexpression in mouse cortical RGs at mid-corticogenesis suppresses ERK signaling and slows the cell cycle

Human cortical fRGs show markedly higher PKA signaling than their mouse counterparts (Fig. 1 and *SI Appendix,* Fig. S2). To test the functional impact of elevated PKA signaling in vivo, we overexpressed constitutively active PKA catalytic subunit mutants—*PRKACA-L206R* and *PRKACA-W196G*—in mouse cortical RGs via in utero electroporation (IUE) and neonatal electroporation. These two mutations, originally identified in adrenal Cushing’s syndrome, generate hyperactive PRKACA variants that drive constitutive PKA signaling (65, 66).

We performed scRNA-Seq analysis of E17.0 mouse cortices. Control littermates received FlashTag at E16.0, whereas experimental littermates received *PRKACA-L206R-GFP* via IUE. Labeled cortical cells were FACS-sorted 24 h later and processed for scRNA-Seq (**Fig. 4*A, B***). Unsupervised clustering of the transcriptomic data revealed that IUE-transduced cortical RGs could be resolved into three distinct subpopulations expressing moderate (RG-M), high (RG-H), and very high (RG-VH) levels of *PRKACA-L206R-GFP* (**Fig. 4*B***), whereas control RGs (GFP-negative) expressed basal levels of endogenous PRKACA typical of normal mouse development (*SI Appendix,* Fig. S20).

**Fig. 4.**
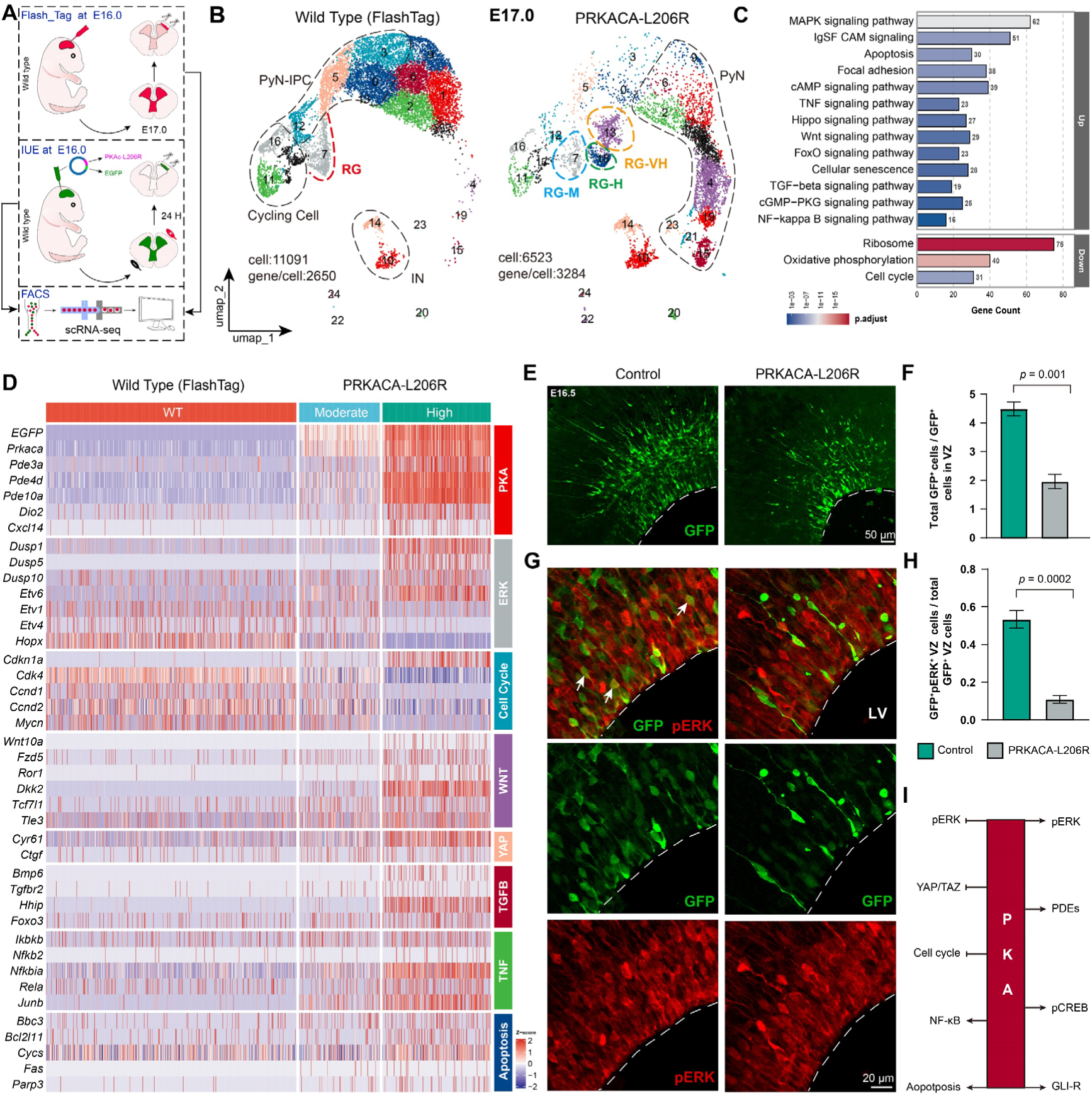
PKA overexpression in mouse cortical RGs at mid-corticogenesis suppresses ERK signaling and slows the cell cycle. (*A, B*) scRNA-Seq was performed on FACS-sorted cortical cells from E17.0 mouse littermates. Controls received FlashTag at E16.0, while experimental embryos were electroporated in utero (IUE) with *PRKACA-L206R-GFP*. Cells were harvested 24 h later. PRKACA-L206R-GFP-positive RGs resolved into three clusters based on GFP intensity: RG-M (moderate, cluster 7), RG-H (high, cluster 18), and RG-VH (very high, cluster 13). (***C***) KEGG enrichment analysis was performed on DEGs comparing PRKACA-L206R-GFP-labeled RG-H to control RGs (WT cluster 7). (***D***) Heatmap analysis of DEGs in RG-M and RG-H versus control RGs revealed dose-dependent PKA effects: suppression of ERK and cell cycle, and enhancement of WNT, apoptosis, TGFB, and TNF signaling. (***E, F***) E14.5 IUE with *GFP* or *PRKACA-L206R-GFP*; analyzed at E16.5. GFP labeling: mutant RGs yield far fewer progeny than controls, reflecting profound cell cycle slowing. (***G, H***) GFP/pERK co-staining: pERK⁺ cells are markedly reduced in *PRKACA-L206R-GFP*-labeled cortical RGs compared with control *GFP*-labeled ones (arrows). (***I***) Schematic summary of the functional effects of PKA hyperactivation in mouse cortical RGs.

Our scRNA-seq analysis identified 3,300 DEGs between *PRKACA-L206R-* expressing RG-H cells and control RGs, with up- and down-regulated genes roughly equally represented. KEGG pathway enrichment analysis of the upregulated genes revealed their involvement in multiple signaling pathways, including MAPK (ERK), cAMP-PKA, apoptosis, growth hormone synthesis/secretion/action, TNF/NF-κB, Hippo, WNT (canonical and non-canonical), FoxO, cellular senescence, TGF-β, and cGMP-PKG signaling. In contrast, the downregulated genes were primarily enriched in cell-cycle-related pathways (**Fig. 4*C***).

Heatmap and feature plot analyses of DEGs in *PRKACA-L206R*-expressing RG-H and RG-M cells (relative to control RGs) confirmed dose-dependent PKA effects, including suppressed ERK signaling, slowed cell cycle, and enhanced activities of PKA, apoptosis, WNT, TGF-β, and TNF pathways (**Fig. 4*D*** and *SI Appendix,* Fig. S20). Elevated PKA signaling strongly induced the upregulation of its negative feedback regulators (*Pde3a, Pde4d, Pde10a*) and direct readout genes (*Dio2* and *Cxcl14*) in cortical RGs (**Fig. 4*D*** and *SI Appendix,* Fig. S20) (21, 22). Notably, high PKA levels also upregulated TGF-β signaling, and TGF-β is known to induce *Hhip* expression (**Fig. 4*D*** and *SI Appendix,* Fig. S20) (67), which potently blocks SHH-SMO signaling. This reveals a novel dual mechanism by which PKA suppresses SHH-SMO activity: directly through PKA-mediated inhibition (68), and indirectly via the TGF-β–HHIP axis. Thus, PKA acts as a broad negative regulator of SHH–SMO signaling.

Notably, although KEGG pathway enrichment analysis of the upregulated genes implicated the MAPK signaling pathway, closer examination of bona fide ERK readout genes—including upregulated *Dusp1/5/10* and *Etv6*, alongside downregulated *Etv1/4* and *Hopx*—revealed that ERK signaling was significantly diminished (**Fig. 4*D*** and *SI Appendix,* Fig. S20). This underscores that KEGG analysis merely indicates which pathways are engaged in a given biological process; by contrast, only genes that are unequivocally established as direct pathway readouts can reliably report whether a given signaling pathway is activated or suppressed.

We next validated these scRNA-Seq findings by immunohistochemistry and IUE in vivo. At E14.5, control littermates received control GFP plasmids, whereas experimental littermates received *PRKACA-L206R-GFP* via IUE. Embryos were harvested 48 h post-electroporation at E16.5. Immunostaining for GFP, pERK, and HOPX revealed that *PRKACA-L206R-GFP*-labeled cortical RGs generated very few IPC progeny, indicating that elevated PKA signaling potently promotes RG quiescence and reduces proliferation (**Fig. 4*E, F***). Moreover, pERK and HOPX expression levels were both significantly reduced in mutant RGs compared with controls (**Fig. 4*G, H*** and *SI Appendix, Fig. S21*). Collectively, these results support the conclusion that PKA overexpression in mouse cortical RGs suppresses ERK signaling, slows the cell cycle, and exerts broad pleiotropic effects (**Fig. 4*I***) (69).

### PKA overexpression in neonatal mouse cortical RGs suppresses SHH–SMO and YAP/TAZ signaling

To assess the effect of elevated PKA signaling in neonatal mouse cortical RGs—at a stage when SHH–SMO and YAP/TAZ are highly active—we overexpressed *PRKACA-W196G-GFP* (another constitutively active PKA mutant) via electroporation at P0. Control littermates received FlashTag, while experimental littermates received *PRKACA-W196G-GFP*. Transduced cortical cells were FACS-sorted 48 h later for scRNA-Seq. Unsupervised clustering revealed three transgene-expressing RG subpopulations (moderate, high, and very high), whereas control RGs (FlashTag-labeled, GFP-negative) maintained basal PRKACA levels typical of P2 (*SI Appendix,* Fig. S22). KEGG enrichment of upregulated genes pointed primarily to cAMP signaling, whereas downregulated genes were enriched in Integrin, Gap junction, cGMP-PKG, Ras, and Hedgehog signaling (*SI Appendix,* Fig. S22). Notably, KEGG analysis did not capture cilia-related pathways. We therefore performed GO term analysis on the downregulated gene set, which revealed significant enrichment in neural precursor cell proliferation, smoothened (SHH–SMO) signaling, and multiple cilia-associated terms—including cilium organization/assembly, ciliary transition zone, basal body, ciliary plasm, photoreceptor connecting cilium, non-motile cilium assembly, and intraciliary transport (*SI Appendix,* Fig. S22). This coordinated downregulation suggests that ciliogenesis, encompassing both primary and multiple cilia, is profoundly suppressed.

Heatmap analysis of dysregulated genes in *PRKACA-W196G* labeled RG-M, RG-H, and RG-VH confirmed dose-dependent PKA effects: suppression of ERK, SHH– SMO, and cell cycle progression, global downregulation of ciliary genes, and enhancement of PKA and apoptosis (*SI Appendix,* Fig. S23). These results are highly consistent with our *PRKACA-L206R* dataset, with the P2 timepoint further highlighting PKA-mediated repression of SHH–SMO and ciliogenesis, driven by potent suppression of YAP/TAZ (*SI Appendix,* Fig. S23).

We validated these findings in vivo by electroporation and immunohistochemistry. At P0, control *Ai9* (tdTomato-tdT reporter) littermates received *Cre* plasmids, whereas experimental littermates received *Cre* together with *PRKACA-L206R-GFP*. Tissues were harvested at 4, 6, 10, and 30 days post-electroporation (controls) and at 4 and 6 days (experimental). Cortical RGs labeled via *Cre* electroporation at P0 retained their full-span morphology until P10, and their differentiated progeny—astrocytes, oligodendrocytes, and ependymal cells—were detected at P30 (*SI Appendix,* Fig. S24). These results confirm that this IUE strategy efficiently labels P0 RGs and their long-term descendants. We next performed double-immunostaining for tdT and FOXJ1, which revealed a marked reduction in tdT⁺/FOXJ1⁺ cells within the cortical VZ of *PRKACA-L206R*-expressing samples relative to controls at both P4 and P6. These findings confirm that elevated PKA activity blocks ependymal differentiation and disrupts multiciliogenesis, consistent with PKA-mediated repression of YAP/TAZ (*SI Appendix,* Fig. S25, S26).

Together, our findings demonstrate that PKA signaling in cortical RGs exerts broad and multifaceted effects. High PKA activity suppresses ERK, cell-cycle progression, SHH–SMO, YAP/TAZ, ciliogenesis, and ependymal cell formation (**Fig. 4*I***). Of note, PKA can either promote or inhibit ERK in a context-dependent manner, as has been well documented in the literature (69–73).

## Discussion

This study presents six main findings: 1) Human cortical fRGs highly express conserved ERK-boosting genes that are low or absent in mice, leading to elevated ERK signaling. 2) Human oRGs show higher ERK/PKA signaling than tRGs across GW16–GW26 (≥10 weeks), ensuring neurogenesis over gliogenesis. 3) *JUNB* does not bidirectionally promote cortical neurogenesis, as it is expressed predominantly in gliogenic RGs in both mice and humans. 4) ERK signaling exerts broad and profound effects in cortical RGs. Overexpressing ERK in mouse cortical RGs upregulates cAMP-PKA, promotes self-renewal, expands the RG pool, and accelerates the early cell cycle. 5) PKA signaling serves a critical function in cortical RGs; its overexpression at mid-corticogenesis represses ERK and attenuates cell-cycle progression. 6) PKA overexpression in neonatal cortical RGs inhibits both SHH–SMO and YAP/TAZ signaling, blocking ependymal cell generation. Based on these and our other studies, we conclude that elevated ERK activity is the most fundamental and core driver of mammalian cortical expansion during development and evolution— providing a coherent, contradiction-free explanation for all major evolutionary changes in cortical RGs (**Fig. 5**).

**Fig. 5.**
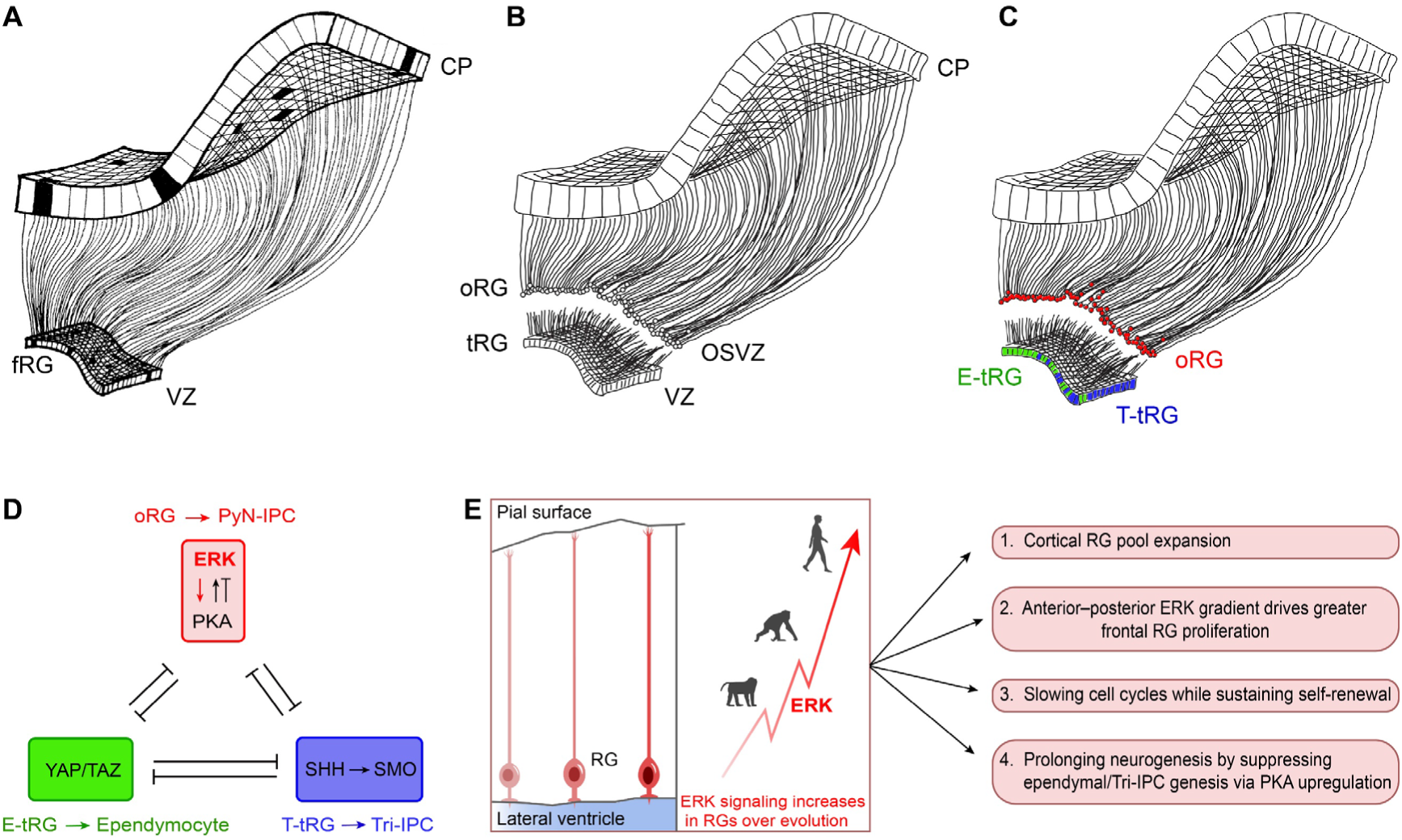
ERK drives four conserved RG hallmarks across evolution to expand the cortex. (***A***) In the human cortex, full-span radial glia (fRGs) initially generate PyN-IPCs committed to deep-layer pyramidal neurons, which migrate to the cortical plate (CP). At this early stage, the scaffold consists of continuous fibers spanning the entire thickness of the developing cortex. (***B***) At the mid-neurogenesis transition (∼GW16), human cortical fRGs undergo major scaffold remodeling: the once-continuous fRG architecture becomes physically discontinuous, giving rise to outer radial glia (oRGs) in the outer subventricular zone (OSVZ) and truncated radial glia (tRGs) in the ventricular zone (VZ). (***C***) tRGs are now recognized as comprising two distinct subtypes: ependymal-generating tRGs (E-tRGs) and Tri-IPC-generating tRGs (T-tRGs). Essentially, during the GW16–GW26 stage, three lineage programs proceed in parallel: oRGs within the OSVZ generate upper-layer PyN-IPCs; E-tRGs initiate ependymal cell differentiation in the medial VZ and progressively expand this program into the lateral VZ; and T-tRGs give rise to Tri-IPCs across the ventrolateral–dorsal VZ. Panels ***A–C*** are modified and adapted from previous studies with permission (15, 54). (***D***) Schematic diagram of the cortical radial glia tri-lineage principle. Cortical RG subtype identity and fate are governed by three pathways: ERK/PKA in neurogenic RGs, YAP/TAZ in ependymogenic RGs, and progressively increasing SHH–SMO in Tri-IPC-generating RGs. These pathways form a cross-repressive network coordinating neurogenesis and gliogenesis. (***E***) From rodents to primates and humans, cortical RG ERK signaling progressively increases over evolution, reflected by deepening red color in the left panel. Concurrently, cortical RGs across species display four conserved hallmarks that collectively promote pyramidal neuron production and drive cortical expansion (right panel). We propose that elevated ERK signaling acts as the common upstream driver of all four hallmarks during evolution.

### The cortical radial glia tri-lineage principle

In this study, we formally propose the cortical radial glia tri-lineage principle, which defines the molecular identity, lineage output, and spatiotemporal dynamics of three cortical RG subtypes. As introduced earlier, human cortical fRGs are neurogenic and generate deep-layer PyNs (**Fig. 5*A***). Following the emergence of tRGs and oRGs around GW16 (**Fig. 5*B***), oRGs sustain the neurogenic function of fRGs and predominantly generate upper-layer PyN-IPCs. Concurrently, tRGs undergo a gliogenic switch: medial E-tRGs produce ependymal cells, whereas ventrolateral– dorsal T-tRGs generate Tri-IPCs (**Fig. 5*C***) (21, 22). At the molecular level, cortical RG subtype identity is governed by a cross-repressive signaling network. ERK/PKA activity drives neurogenesis while suppressing YAP/TAZ and SHH–SMO; YAP/TAZ promotes ependymal differentiation while inhibiting the other two pathways; and progressively increasing SHH–SMO specifies Tri-IPC fate by repressing PKA and weakly suppressing ERK (**Fig. 5*D***). Thus, these three pathways—ERK/PKA, YAP/TAZ, and SHH–SMO—coordinate the balance between neurogenesis and gliogenesis across cortical regions and species (21, 22).

Cross-species comparisons reveal that this tri-lineage logic is evolutionarily conserved but temporally regulated differently. In mice, the three lineage programs unfold sequentially along a spatial gradient: N-RGs first generate PyN-IPCs, followed by medial E-RGs and ventrolateral–dorsal T-RGs (21, 22). Importantly, most mammalian species possess a gyrencephalic cortex and therefore likely all harbor oRGs, indicating that this principle represents a fundamental feature of cortical RG progression across the vast majority of mammals. Mice and rats are evolutionary outliers in this regard. Although lissencephalic, they originated from a larger, gyrencephalic ancestor—as evidenced by the fact that gyrencephalic ferrets diverged from the human lineage before rodents did (4, 74). During evolution, under pressures from increasing litter size and decreasing body and brain size, mice and rats likely lost oRGs and retained only full-span RGs, each producing merely 8–9 PyNs to complete cortical development rapidly (75). In contrast, human cortical neurogenesis—mediated by both fRGs and oRGs—extends for at least 18 weeks (GW8–GW26), with each fRG (and its oRG progeny) generating 60–90 PyNs (4), thereby supporting the massive expansion of the human neocortex.

### High ERK/PKA signaling enables sustained oRG neurogenesis via a unique mechanism

High ERK signaling is well established as a promoter of rapid cell division, as observed in tumor cells, glial IPCs (e.g., via EGFR–ERK), and many other actively dividing cell types (76). In human fRGs and oRGs, however, elevated ERK signaling does not accelerate cell cycle progression. Instead, evolution has driven progressively slower cell cycles while preserving self-renewal. Consequently, human oRGs may have the longest cell cycle in the human brain, lasting approximately 50– 60 hours or more, compared to ∼18–24 hours in mouse cortical RGs and ∼30–40 hours in macaques (4, 74, 77).

Relatively higher ERK signaling enables prolonged neurogenesis in human fRGs and oRGs through a unique molecular mechanism (**Fig. 5*E***). Human oRGs exhibit concomitantly elevated ERK and PKA signaling, which precludes SHH–SMO activity—notably, SHH is a potent morphogen and mitogen. This specific signaling environment (or niche) thus uncouples high ERK/PKA activity from accelerated cell division. Instead, without the mitogenic function of SHH-SMO, elevated GLI-repressor levels, together with ERK-driven upregulation of BMP7 and HOPX, as well as elevated PDGFD/PDGFRB and CXCL14/CXCR4 levels—all known to promote cell cycle arrest—collectively contribute to the exceptionally long cell cycle observed in oRGs. Importantly, despite their protracted cell cycle, human oRGs retain classic stem cell features and do not enter permanent terminal quiescence, owing to elevated ERK and LIFR signaling—both well established to sustain self-renewal.

Longer cell cycles promote genomic maintenance, lower mutation rates, fate regulation, and lineage fidelity. This extended timeframe gives oRGs more opportunity to respond to growth factor and transcription factor cues and to prepare for asymmetric division—facilitating robust partitioning of organelles, mRNAs, and proteins into daughter cells, thereby generating distinct progeny fates (one self-renewing, the other neuronal). These features enable human oRGs to sustain prolonged neurogenesis, generate upper-layer PyNs with precision, and maintain self-renewal without gliogenic switching.

While others have proposed that cortical oRGs generate cortical interneurons, OPCs, or Tri-IPCs (28, 37, 40, 78–82), and that glioblastoma may harbor oRG-like cells with cancer-like stem cell properties (83), our findings indicate that oRGs predominantly produce PyNs. Although SHH-SMO signaling has been proposed to drive cortical expansion (84–89), our functional studies indicate that its primary role is to promote Tri-IPC generation in both humans and mice, rather than directly driving expansion (4, 16, 19, 21, 22, 27, 29–31). Based on our analysis, these alternative interpretations seem difficult to reconcile with the available evidence and may therefore be less likely.

### ERK-driven cortical expansion in evolution: a synthesis of established evidence

The emerging picture of ERK-driven cortical expansion in evolution is increasingly coherent. Three pathways—ERK/PKA, YAP/TAZ, and SHH–SMO—constitute a tripartite regulatory network defined by cross-repressive interactions (**Fig. 5*D***) (21, 22). This network likely originated as early as ∼500 million years ago, based on co-expression of *Fgf8* and hedgehog homologs in the ectoderm of the hemichordate *Saccoglossus kowalevskii* (90). The presence of hedgehog signaling in this ancient organism implies concurrent cAMP–PKA activity, itself an ancient GPCR-mediated mechanism. The Hippo–YAP/TAZ pathway, a mechanosensory module that responds to cell contact, mechanical tension, and spatial crowding, also emerged early— potentially in the hemichordate lineage or even earlier in multicellular forebears (72).

Fast-forward to vertebrates—from fish and amphibians to reptiles, birds, and mammals—all possess a pallium (cortex) (91), indicating the presence of cortical RGs and associated ERK and cAMP–PKA signaling. This signaling axis safeguards pallial neurogenesis while suppressing ependymal and glial fates through inhibition of YAP/TAZ and SHH–SMO. Across the evolutionary trajectory from vertebrates to primates and humans, cortical RGs exhibit four conserved hallmarks: (1) expansion of the RG pool; (2) frontal RG proliferation driving frontal cortex growth; (3) slower cell cycles with sustained self-renewal; and (4) prolonged neurogenesis (**Fig. 5*E***).

Collectively, these features boost pyramidal neuron output and cortical size. We propose elevated ERK signaling as the unifying driver. ERK expands the RG pool and sustains self-renewal (21, 31). FGFs at the anterior neural ridge promote frontal telencephalic growth via ERK (92). ERK also upregulates PKA, BMP7, HOPX, and other factors to lengthen the cell cycle without compromising self-renewal (21, 31). Within the tri-lineage principle framework, ERK prolongs neurogenesis via PKA while repressing gliogenesis (**Fig. 5*E***).

Thus, even a modest increase in developmental ERK activity within cortical RGs suffices to drive cortical expansion. As François Jacob famously observed, evolution is a tinkerer—repurposing existing components for new functions or assembling them into greater complexity (93). This perspective posits that a conserved genetic and biochemical machinery—which we identify as ERK signaling (Fig. 5)—underlies the evolutionary increase in cortical neuron numbers. Certainly, other factors may converge on or complement this pathway, yet their collective actions likely funnel into progressively heightened ERK activity in cortical RGs over evolutionary time, ultimately driving cortical expansion and brain enlargement.

## Materials and Methods

### A full description of materials and methods is available in *SI Appendix*

All experiments were performed in accordance with the guidelines of Shanghai Medical College of Fudan University and were approved by the Institutional Animal Care Committee (20240229-193 and YSA20260135). *SI Appendix* contains additional details regarding the mouse lines, mouse handling and sample processing, and human cerebral cortex sample collection.

For scRNA-seq, *in utero* electroporation (IUE) labeling, or FlashTag labeling followed by FACS were used to enrich late cortical progenitor cells and their progeny from transgenic mice and their respective controls. Detailed information is provided in the online *SI Appendix*.

Immunohistochemistry was performed using a regular protocol. Primary antibodies were diluted in blocking solution and incubated overnight at 4°C. Sections were incubated with the indicated secondary antibodies for 2 h at room temperature. Details regarding primary antibodies, dilutions, imaging, and analysis are described in *SI Appendix*.

Plasmid construction, scRNA-seq and data analysis, IUE, *in situ* hybridization, image acquisition and analysis, quantification and statistical analysis were performed according to published protocols. Information for detailed experimental procedures and analyses can be found in online *SI Appendix*.

### Data, Materials, and Software Availability

In this study, we generated 7 scRNA-Seq samples from mouse cortical tissue, with FACS enrichment applied where indicated. The samples included: (1) E12.0 whole control cortex; (2) E12.0 whole *Emx1-Cre; Rosa^MEK1DD/+^* cortex; (3) E17.0 control cortex, which received FlashTag-CellTrace Yellow injection at E16.0 and was FACS-sorted at E17.0; (4) E17.0 cortex overexpressing *PRKACA-L206R-GFP* via IUE at E16.0 and FACS-sorted at E17.0; (5) P2 control cortex, with FlashTag injection at P0 and FACS sorting at P2; (6) P2 cortex overexpressing *PRKACA-W196G-GFP* via IUE at P0 and FACS-sorted at P2; and (7) P0 WT cortex labeled with FlashTag at E18.0 and FACS-sorted at P0. All scRNA-seq data have been deposited in the Gene Expression Omnibus (GEO) under the accession number **GSE341151**. Additional scRNA-seq datasets were obtained from published studies; further details are available in online *SI Appendix*.

## Acknowledgments

This study was supported by the Ministry of Science and Technology of China (STI2030-2021ZD0202300) and National Natural Science Foundation of China (NSFC 32100768, 32200776, and 32200792).

## Author Contributions

Z.Y. designed research. T.F, Z.Z, and W.Z. performed research. T.F., D.Q., and Z.Y. contributed new reagents/analytic tools; C.Y., Z.S., D.H., J.L., F.Y., J.Y., Z.L., Y.Y., T.M., W.L., G.L., X.S., D.Q., D.M., W.H., and Z.X. analyzed data. Z.Y. wrote the paper.

