## supplemental files for "ERK signaling integrates multiple mechanisms to drive cortical evolutionary expansion"

**This PDF file includes:**

Supporting text

Figures S1 to S26

SI References

### Supporting Information Text

### Materials and Methods

All experiments were performed in accordance with the guidelines of Shanghai Medical College of Fudan University and were approved by the Institutional Animal Care Committee (20240229-193 and YSA20260135). Histology, transgenic assays, plasmid construction, *in utero* electroporation (IUE), mouse mRNA *in situ* hybridization, scRNA-Seq analysis, microscopy and imaging, and quantification and statistical analysis were performed and analyzed according to published protocols.

### Mice

All animal procedures were approved by the Animal Ethics Committee of Fudan University and performed in accordance with institutional guidelines. *Emx1-Cre* (JAX no. 005628) (1), *hGFAP-Cre* (JAX no. 004600) (2), *Rosa*$^{MEK1DD/+}$ (JAX no. 012352) (3), and *Ai9* (JAX no. 007909) (4), were described previously. *Map2k1 (exon 2) flox* and *Map2k2 (exon 4-9) flox* mice (5), and *Egfr* flox mice (6), were purchased from GemPharmatech Co., Ltd., Nanjing, China. The presence of a vaginal plug was defined as embryonic day 0.5 (E0.5), while the day of birth was defined as postnatal day 0 (P0). Sexes were not determined for embryonic and early postnatal animals. For immunofluorescence analyses, brain samples were routinely collected from three independent experiments unless otherwise indicated (n values represent the number of individual brains analyzed).

The genotypes of the control samples are provided below. Unless otherwise noted, all controls were littermates of the corresponding experimental groups. For the *Emx1-Cre; Rosa*$^{MEK1DD/+}$ experimental mice, controls were either *Emx1-Cre* without *Rosa*$^{MEK1DD/+}$ or *Rosa*$^{MEK1DD/+}$ without *Emx1-Cre*. For the *hGFAP-Cre; Egfr* conditional knockout (cko) experimental mice, controls were either *hGFAP-Cre* without *Egfr*-flox or *Egfr*-flox without *hGFAP-Cre*. For *Emx1-Cre; Map2k1/2* double conditional knockout (dcko) embryos, controls were *Map2k1* and *Map2k2* flox littermates lacking *Emx1-Cre*.

### Brain tissue preparation

Mouse embryos were separated from deeply anesthetized pregnant mice. Brain were dissected out and fixed in 4% paraformaldehyde (PFA) pre-treated with diethylpyrocarbonate overnight. Postnatal mice were deeply anesthetized and transcardially perfused with phosphate-buffered saline followed by 4% PFA. All brains were post-fixed overnight in 4% PFA at 4℃ and dehydrated in 30% sucrose for at least 24 hours. Then the brains were embedded with O.C.T. (Sakura Finetek) in ethanol slush with dry ice and stored at -80℃. Mouse brains in this study were sectioned into 20 μm or 40 μm.

The GW19 and GW23 human cortical sections with high quality were of 60 μm. The specimen originally assigned to GW18 was re-evaluated comprehensively using immunohistochemical criteria and was accordingly re-designated as GW19 (5, 7-12). All sections were mounted on glass slides for staining; their characterization has been described in our earlier studies (5, 7-12).

In this study, we collected a new human brain sample at GW16 under strict ethical compliance. The human fetal brain tissue was obtained from Shanghai Changning District Maternal and Child Health Hospital following elective termination of pregnancy, with written informed consent provided by the donors. The decision to terminate the pregnancy was made independently of this study. All procedures were approved by the Institutional Review Board of Shanghai Changning District Maternal and Child Health Hospital (Approval No. CNFBLLKT-2025-007) and were conducted in accordance with relevant ethical guidelines. The sample was de-identified prior to research use.

#### Plasmid construction

To generate the mouse cDNA equivalent of the human PRKACA-L206R mutation, we introduced two nucleotide substitutions into the full-length wild-type mouse *Prkaca* cDNA, converting the leucine codon (TTG) at position 206 to an arginine codon (CGG). To generate the mouse cDNA equivalent of the human *PRKACA*-W196G mutation, we introduced a point mutation into the full-length wild-type mouse *Prkaca* cDNA, converting the tryptophan codon (TGG) at position 196 to a glycine codon (GGG).

The pCAG-ires-GFP backbone (Addgene #11150) was used for cloning. cDNA fragments encoding the mouse equivalents of the human *PRKACA-L206R* and *PRKACA-W196G* mutations were inserted into this vector to create the respective overexpression constructs, designated *PRKACA-L206R-GFP* and *PRKACA-W196G-GFP*.

#### *In Utero* electroporation

*In utero* electroporation (IUE) was performed as described previously (13). Briefly, pregnant mice were anesthetized with isoflurane using an animal anesthesia machine. Plasmid solution (1–2 μg/μl, 0.5 μl per embryo) mixed with 0.05% Fast Green (Sigma) was injected into the lateral ventricle of each embryo through a beveled glass micropipette. Five electrical pulses (50 ms duration) were delivered across the uterine wall at optimized voltages using 7-mm platinum tweezertrodes (BTX, 45-0488) connected to an electroporator (BTX, ECM830), with a 950-ms interval between pulses. Voltages were adjusted according to embryonic age: 34 V for E13, 36 V for E15, and 37 V for E16. Electroporated brains were harvested and analyzed at time points specified in the main text.

Electroporation of neonatal mice was performed similarly to IUE, with the following modifications. Briefly, plasmid solution (1–2 μg/μl, 1.0 μl per pup) mixed with 0.05% Fast Green (Sigma) was injected into the lateral ventricle of each neonatal mouse through a beveled glass micropipette. Five electrical pulses (50 ms duration, 950-ms intervals) were delivered at 100 V using 7-mm platinum tweezertrodes (BTX) placed directly on the neonatal skull, connected to an electroporator (BTX, ECM830).

### ***In situ* hybridization**

*In situ* hybridization assays were performed using digoxigenin-labeled riboprobes on 20-μm frozen sections as previously described (13). DNA templates for *in situ* hybridization probes were obtained by PCR from mouse cDNA library. DNA fragments obtained by PCR were cloned into *pBluescript II KS (*-) vectors. The probes with a final concentration of 0.5 ng/μl were used for hybridization.

The following primers were used for generating the DNA templates:

*Etv5*-F: TTCATGGCCTCCCACTGAAAATC
*Etv5*-R: CCTTCGTTGTAGGGGTGAGGGTT

*Spry2*-F: CAGATGTGTTCTAAGCCTGCTG
*Spry2-R: TCCCATAGTCAATGACGTTCTG*

### **Immunohistochemistry**

Immunohistochemical staining was carried out using standard protocols on coronal sections from mouse, macaque, and human brains. Sections of 20-μm (mouse), 30-μm (E55 macaque), and 60-μm (human) were stained on glass slides, whereas 40-μm mouse sections were processed as free-floating sections.

Sections were rinsed with TBS (0.01 M Tris–HCL + 0.9% NaCl, pH7.4) for 10 minutes (mins), incubated in 0.5% Triton-X-100 in TBS for 30 mins at room temperature (RT), and then incubated in a blocking buffer (3% donkey serum + 0.5% Triton-X-100, pH7.2) in TBS for 2 hours. After blocking, the sections were incubated with primary antibodies (diluted in the blocking buffer) for overnight at 4°C. On the next day, the sections were washed in TBS three times for 10 mins each, and incubated with secondary antibodies (1:500, all from Jackson ImmunoResearch) for 2 hours in dark at RT. Then they were rinsed three times with TBS for 10 mins each and incubated with 4',6-diamidino-2-phenylindole (DAPI) (Sigma, 400 ng/mL) for 2 mins.

The following primary antibodies were used in this study: chicken anti-GFP (1:3000, Aves labs, GFP-1020), rabbit anti-pERK1/2 (pMAPK, 1:400, Cell Signaling Technology, #4370), rabbit anti-SOX9 (1:1000, Abcam, ab185966), goat anti-SP8 (1:3000, Santa Cruz, Sc-104661 ), rabbit anti-HOPX (1:500, Proteintech, 11419-1-AP), goat anti tdT (tdTomato, 1:2000, SICGEN, Ab8181), rabbit anti SOX2 (1:1000, Abcam, ab97959), at anti-GFAP monoclonal antibody (1:1000, Invitrogen, Cat# 13-0300), goat anti-EGFR (1:1000, R&D System, BAF1280), rabbit anti-OLIG2 (1:500, Millipore, AB9610), rat anti-OLIG2 (1:500, Oasis Biofarm, OB-PRB009-01), and mouse anti FOXJ1 (1:1000, Invitrogen Cat# 14-9965-80).

**Image acquisition**

Whole-slide imaging (WSI) was performed utilizing an Olympus VS200 automated slide scanner configured with high-numerical-aperture 10× and 20× objectives, enabling high-throughput visualization of full tissue sections. For detailed subcellular characterization, high-resolution fluorescence imaging was conducted on an Olympus FV3000 confocal microscope system equipped with advanced 20× and 40× objectives, optimized for precise multi-channel signal detection. To ensure publication-quality visualization, all images were systematically processed (merged, cropped, and contrast-adjusted) using Adobe Photoshop and Illustrator software. These post-acquisition adjustments were applied uniformly across all experimental groups, focusing strictly on linear scaling of brightness and contrast while maintaining absolute data integrity without any selective pixel manipulation or signal enhancement.

**scRNA-Seq**

Whole cortices from E12 and E15 mouse embryos were dissected, dissociated into single cells, and subjected to scRNA-seq. For cortical samples from mice older than E15.0, two labeling strategies were employed prior to scRNA-seq.

For FlashTag labeling, 0.5 μl of 10 mM CellTrace Yellow (Life Technologies, #C34567) was injected into the lateral ventricles. After 24–72 hours, the brains were immediately removed and submerged in fresh ice-cold Hanks' balanced salt solution (Gibco 14175-095). The dorsal cortices were then dissected, minced, and dissociated into single-cell suspensions using a Papain Cell Dissociation Kit according to the manufacturer's instructions. FlashTag-positive cells were enriched by fluorescence-activated cell sorting (FACS).

Alternatively, for the two scRNA-seq experiments involving PKA overexpression, E17.0 cortices were IUE with *PRKACA-L206R-GFP* at E16.0 and FACS-sorted at E17.0; P2 cortices were electroporated with *PRKACA-W196G-GFP* at P0 and FACS-sorted at P2. In both cases, the yield of GFP-positive cells was lower than anticipated. To obtain sufficient cells for downstream library preparation, we therefore adjusted the FACS gating to be less stringent to accommodate the lower yield. Importantly, nearly all cortical RGs included in the final dataset were GFP-positive, confirming that the transcriptomes were derived exclusively from successfully transduced cells. Notably, many GFP-negative cells—representing other cell types—were also present in the sorted fractions; however, these were excluded from further analysis.

For FlashTag labeling, 0.5 μl of 10 mM CellTrace Yellow (Life Technologies, #C34567) was injected into the lateral ventricles. After 24–72 hours, the brains were immediately removed and submerged in fresh ice-cold Hanks' balanced salt solution (Gibco 14175-095). The dorsal cortices were then dissected, minced, and dissociated into single-cell suspensions using a Papain Cell Dissociation Kit according to the manufacturer's instructions. FlashTag-positive cells were enriched by fluorescence-activated cell sorting (FACS). Alternatively, for the two scRNA-seq experiments, E17.0 cortices were electroporated with *PRKACA-L206R-GFP* at

E16.0 and FACS-sorted at E17.0; P2 cortices were electroporated with *PRKACA-W196G-GFP* at P0 and FACS-sorted at P2. In both cases, the yield of GFP-labeled cells was lower than anticipated following electroporation. To obtain sufficient cell numbers for downstream library preparation, we adjusted the FACS gating strategy to be less stringent. Nevertheless, it is important to note that the cortical RGs analyzed in the final dataset were all GFP-positive, confirming that the transcriptomes were derived exclusively from successfully transduced cells. GFP negative cells were present across all clusters but were excluded from our analysis; their exclusion did not compromise the validity of our conclusions regarding RG biology.

Following FACS, all sorted cells were subsequently processed for high-throughput scRNA-Seq. Single-cell microfluidic encapsulation and subsequent library construction were performed using the Chromium droplet-based sequencing platform (10x Genomics), strictly adhering to the manufacturer’s protocols (User Guide, document part number: CG00052 Rev C). The resulting cDNA libraries underwent fragment-size purification and absolute quantification utilizing an Agilent 2100 Bioanalyzer system to ensure optimal library complexity and quality. Finally, the pooled libraries were multiplexed and sequenced on an Illumina NovaSeq 6000 platform using a paired-end sequencing strategy to achieve the target read depth required for robust transcriptomic profiling.

**1) scRNA-Seq data analysis**

Raw sequencing FASTQ files were processed directly using the Cell Ranger software pipeline (10x Genomics, version 7.0.1). Cell Ranger performed initial read quality filtering, cell barcode identification, and Unique Molecular Identifier (UMI) counting, followed by alignment to the mouse reference genome (mm10, version 1.2.0). Notably, reads mapping to intronic regions were included in the UMI counting. To ensure high-quality data for downstream analysis, further stringent cell-level quality control was implemented. Specifically, genes expressed in fewer than 3 cells were excluded. Cells were retained only if they met the following criteria: a minimum of 500 detected genes and a mitochondrial gene expression fraction of less than 10%. Additionally, anomalous outlier cell clusters exhibiting generally low-quality transcriptomic metrics were identified and removed prior to downstream analyses.

Post-filtering cell numbers and genes-per-cell counts are shown in the figure as part of the scRNA-Seq quality assessment. E12.0 MEK1DD mice cortex (7931 cells, 3773 genes/cell), E12.0 control mice cortex (7563 cells, 3562 genes/cell), E17.0 PRKACA-L206R-IUE mouse cortex (6523 cells, 3284 genes/cell), E17.0 FlashTag-labeled control mouse cortex (11091 cells, 2650 genes/cell), P0 FlashTag-labeled WT-control mouse cortex (13894 cells, 2891 genes/cell), P2 PRKACA-W196G-IUE mouse cortex (8211 cells, 3006 genes/cell), P2 FlashTag-labeled control mouse cortex (10920 cells, 2871 genes/cell). These scRNA-Seq data have been deposited in the Gene Expression Omnibus (GEO) under the accession number GSE341151.

For data normalization, a global-scaling "LogNormalize" method was applied to the raw count matrix. This method normalizes the gene expression measurements for each cell by the total expression, multiplies by a scale factor of 10,000, and natural-log transforms the result. These normalized data were utilized for downstream differential expression analyses. Prior to dimensionality reduction and clustering, the data were scaled, and potential sources of variation driven by cell cycle phases were strictly regressed out.

The scaled z-scored residuals were used for principal component analysis (PCA). Statistically significant principal components determined by a resampling test were kept for uniform manifold approximation and projection (UMAP) analysis.

Differentially expressed genes (DEGs) among clusters were identified by comparing cells in each cluster against all other cells using the Wilcoxon rank sum test. DEGs were generally defined as those with an adjusted p-value < 0.05; in some analyses, an alternative threshold of nominal $p < 1 \times 10^{-4}$ was applied to select highly significant markers. All these analyses were performed in the Seurat package v5.1.0.

**2）Functional Enrichment Analysis**

To elucidate the biological functions and signaling pathways associated with the identified differentially expressed genes (DEGs), Gene Ontology (GO) and Kyoto Encyclopedia of Genes and Genomes (KEGG) pathway enrichment analyses were performed independently for each individual sample within the human and mouse datasets. Initially, DEGs from each sample were filtered based on a significance threshold of $P < 1e\text{-}5$. The filtered genes were then categorized into up-regulated (log2 fold change > 0) and down-regulated (log2 fold change < 0) gene sets. Prior to the analysis, gene symbols were converted to Entrez gene IDs using the appropriate species-specific genome annotation databases based on the sample's origin: org.Hs.eg.db for human samples and org.Mm.eg.db for mouse samples.

Both GO and KEGG enrichment analyses were conducted independently for the up-regulated and down-regulated gene sets of every sample using the clusterProfiler package(version 4.18.4) in R. For the KEGG pathway analysis, the organism parameters were set to 'hsa' for human samples and 'mmu' for mouse samples. The GO enrichment analysis encompassed all three sub-ontologies: Biological Process (BP), Cellular Component (CC), and Molecular Function (MF). By default, the Benjamini-Hochberg (BH) method was applied for multiple testing correction. GO terms and KEGG pathways with an adjusted *P*-value < 0.05 were considered statistically significantly enriched.

**3）Cross-species Orthologous Gene Mapping**

To eliminate discrepancies in interspecies genomic annotations and enable precise transcriptomic comparisons, we constructed a strictly defined cross-species orthologous gene space. Utilizing the biomaRt R package (version 2.66.1), we accessed the Ensembl database to retrieve genomic annotations for human (hsapiens_gene_ensembl), mouse (mmusculus_gene_ensembl), and macaque (mmulatta_gene_ensembl). With human genes serving as the reference, we extracted the corresponding human orthologs for the murine and macaque gene lists, strictly filtering out many-to-one or unmapped ambiguous annotations. Subsequently, we identified the intersection of shared orthologous genes across the raw single-cell expression matrices (counts) of the three species. The raw matrices were then subsetted to this intersection, unifying the human, macaque, and mouse single-cell expression data into a standardized 1:1:1 orthologous feature space for downstream analysis.

**4）Cross-species Integration and Principal Component Analysis (PCA)**

To mitigate sequencing batch effects and inherent species-specific background differences in the multi-species joint analysis, while preserving critical developmental and evolutionary trajectories, we performed global integration on the ortholog-subsetted single-cell data. Initially, all cells from the three species were merged and normalized, followed by the identification of 2,000 highly variable features. During Canonical Correlation Analysis (CCA), to prevent data bias driven by animal models, the human dataset (GW12) was strictly designated as the reference to compute cross-dataset integration anchors. These anchors were applied to perform batch correction and integration across the entire cell population (IntegrateData). Subsequently, the integrated matrix (integrated assay) was scaled and subjected to Principal Component Analysis (PCA), extracting the first 30 principal components (PCs). This global principal component space served as the unified mathematical coordinate system for evaluating cross-species transcriptomic convergence.

**5）Human Nearest-Neighbor (Hu Neighbor) Analysis**

To quantitatively assess the transcriptomic convergence of experimental murine radial glia (RG) toward early human developmental states at single-cell resolution, we designed a nearest-neighbor (NN) analysis approach based on a principal component (PC) space encompassing four distinct biological groups: experimental mouse, control mouse, wild-type macaque, and wild-type human. First, the coordinate subset for cells annotated as RG was precisely extracted from the globally integrated PCA space, ensuring that distance metrics were not confounded by the topological structure of other cell types. Within this purified RG space, Euclidean distances were used to define the local neighborhood for each target cell (selecting the $k$ = 5 true nearest neighbors after excluding the cell itself).

To quantify the degree of human-like transcriptomic signatures, we calculated the proportion of cells within the local neighborhood of each non-human RG cell (i.e., experimental mouse, control mouse, and wild-type macaque) that originated from the wild-type human dataset. This approach generated a "Human Neighbor" probability distribution at the single-cell level. To validate the statistical significance of the convergence effect in the experimental group, a one-sided Wilcoxon rank-sum test was employed to evaluate the differences in Human Neighbor proportions specifically between the experimental and control murine groups. Furthermore, to visually demonstrate the multi-species relationship and the direction of convergence, the RG cell populations from all four groups were projected onto a two-dimensional space defined by the first two principal components (PC1 and PC2). By calculating the 2D mathematical centroids for each of the four biological groups and plotting 95% core population confidence ellipses alongside centroid shift vectors, we physically corroborated the cross-species transcriptomic reprogramming trajectory driven by the experimental perturbation.

**Data and materials availability**

In this study, we generated 7 scRNA-Seq samples from mouse cortical tissue, with FACS enrichment applied where indicated. The samples included: (1) E12.0 whole control cortex; (2) E12.0 whole *Emx1-Cre; Rosa*$^{MEK1DD/+}$ cortex; (3) E17.0 control cortex, which received FlashTag-CellTrace Yellow injection at E16.0 and was FACS-sorted at E17.0; (4) E17.0 cortex overexpressing *PRKACA-L206R-GFP* via IUE at E16.0 and FACS-sorted at E17.0; (5) P2 control cortex, with FlashTag injection at P0 and FACS sorting at P2; (6) P2 cortex overexpressing *PRKACA-W196G-GFP* via IUE at P0 and FACS-sorted at P2; and (7) P0 WT cortex labeled with FlashTag at E18.0 and FACS-sorted at P0. All scRNA-seq data have been deposited in the Gene Expression Omnibus (GEO) under the accession number GSE341151

Additional scRNA-Seq datasets of mouse cortex used here were derived from our previously published studies: E15.0 whole mouse cortex (GSE293205) (14); E17.0 control cortex, P2 control cortex, and P2 *hGFAP-Cre; Egfr-cko* cortex (GSE282783) (6).

The scRNA-seq data for GW12 and GW14 human cortex (15), are available in the Genome Sequence Archive (National Genomics Data Center, China) under accession HRA000858 at https://ngdc.cncb.ac.cn/gsa-human.

The scRNA-seq data for GW17–GW18 human cortex (16), used to generate the t-SNE plots, were retrieved from the CoDEx web portal (http://solo.bmap.ucla.edu/shiny/webapp/).

The scRNA-Seq data for GW18, GW22, GW23, and GW26 human cortex (GSE162170) have been published previously (17).

The scRNA-Seq data of the E62–E64 macaque frontal cortex (GSE226451) were reported previously (18).

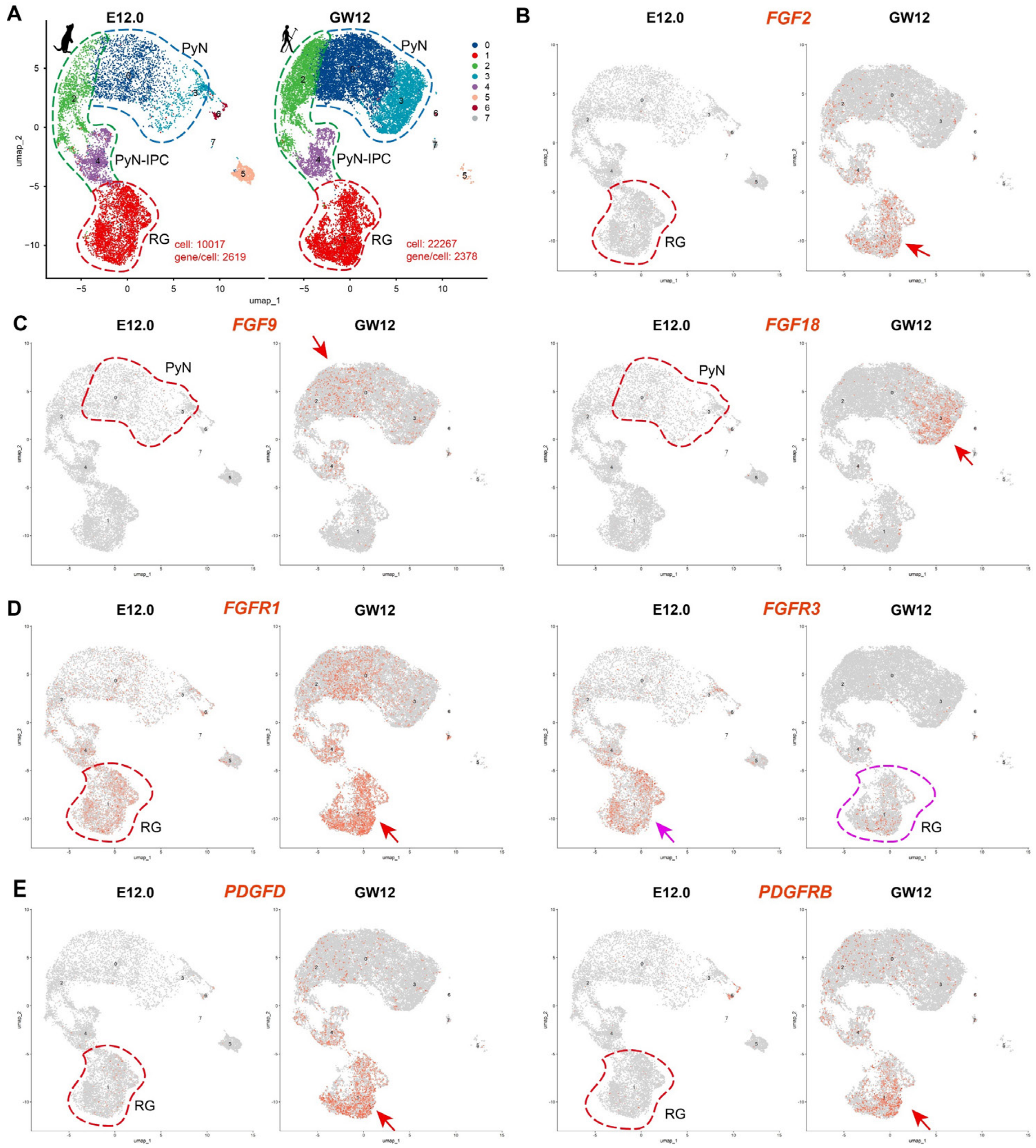


**Fig. S1. Many ERK-boosting genes are highly expressed in human fRGs but low or absent in mice.** (***A***) UMAP projection of human GW12 and mouse E12.0 cortical scRNA-Seq data, colored by cluster. Same as Fig. 1A. (***B***) *FGF2* is readily detected in human cortical fRGs at GW12 but is low or absent in mouse RGs at E12.0. (***C***) *FGF9* and *FGF18* are readily detected in human cortical young deep-layer PyNs at GW12 but are low or absent in mouse counterparts at E12.0. (***D***) *FGFR1* is highly expressed in human cortical fRGs at GW12 but is low in mouse RGs at E12.0. Conversely, *FGFR3* shows the opposite pattern, as ERK signaling suppresses its expression—further confirming that ERK activity is elevated in human cortical RGs at GW12. FGFRs bind FGF2, FGF9, and FGF18 to enhance ERK signaling. (***E***) *PDGFD* and *PDGFRB* are readily detected in human cortical fRGs at GW12 but are low or absent in mouse cortical RGs at E12.0, and this pathway enhances ERK signaling activity. Red arrows (human GW12) and magenta arrow (mouse E12.0) denote high expression.

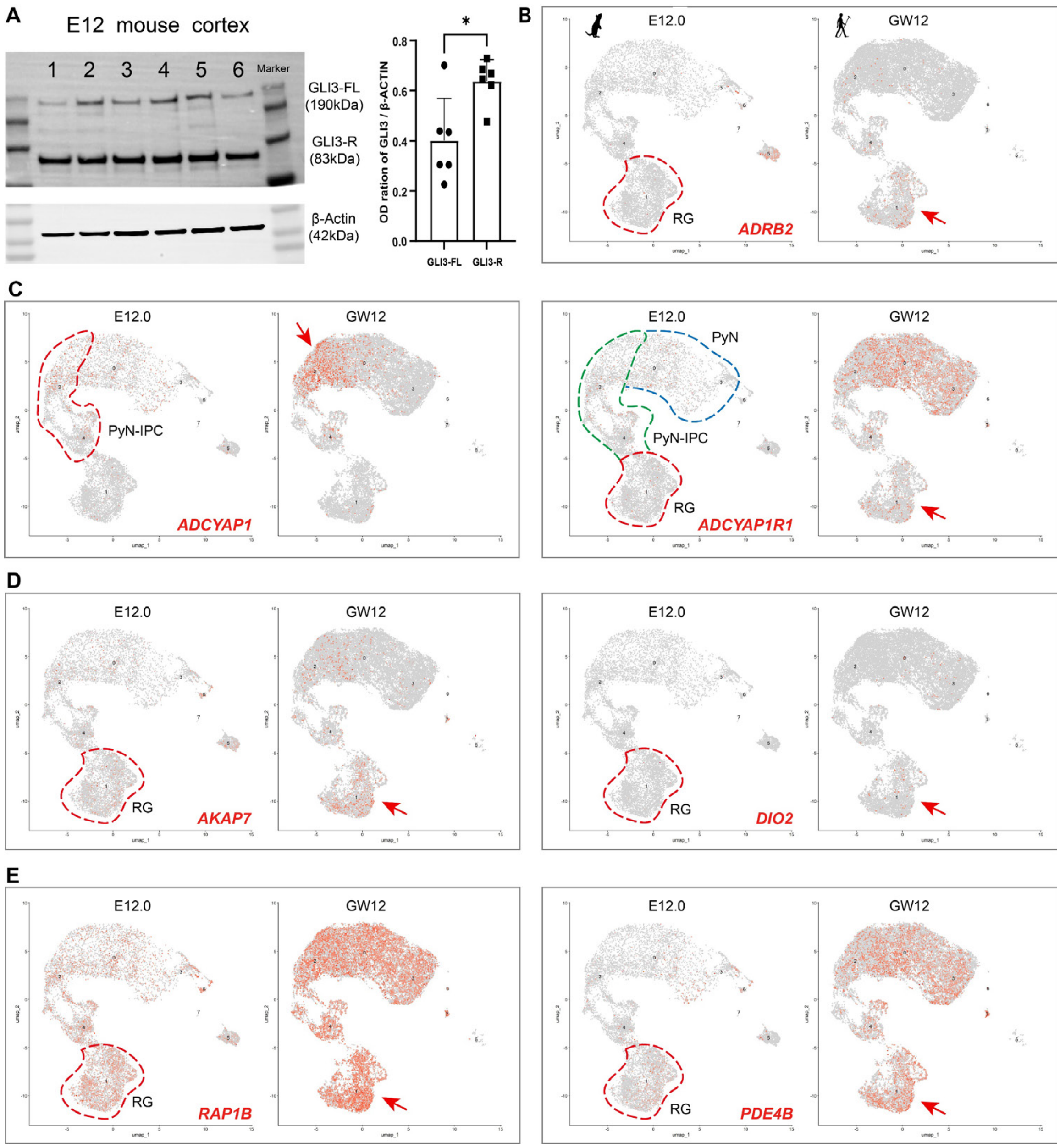


**Fig. S2. cAMP–PKA genes and targets are high in human cortical fRGs but low or absent in mouse RGs.** (***A***) GLI3 western blot of mouse E12 cortex reveals GLI3-R predominance over GLI3-FL, reflecting elevated PKA signaling and enhanced GLI3 cleavage into its repressor form. (***B***) *ADRB2* is present in human cortical fRGs at GW12 but nearly absent in mouse RGs at E12.0, and upon ligand binding, it activates the Gs–AC–cAMP–PKA cascade*.* (***C***) *ADCYAP1* in human PyN-IPCs and *ADCYAP1R1* in fRGs are high at GW12 but low in mouse at E12.0. Their interaction engages Gs–AC–cAMP–PKA. (***D***) *AKAP7*: high in human fRGs (GW12), low in mouse RGs (E12.0); anchors PKA for local cAMP-triggered activation. *DIO2* is upregulated in human GW12 RGs vs. mouse E12 RGs (adj. P = 0.0096) and serves as a PKA signaling readout via direct transcriptional activation through a cAMP-response element. (***E***) *RAP1B* is highly expressed in human GW12 cortical fRGs compared to mouse, consistent with enhanced PKA signaling, whereas *PDE4B* is also upregulated, likely as a negative feedback response to elevated PKA signaling. Red arrows indicate high expression.

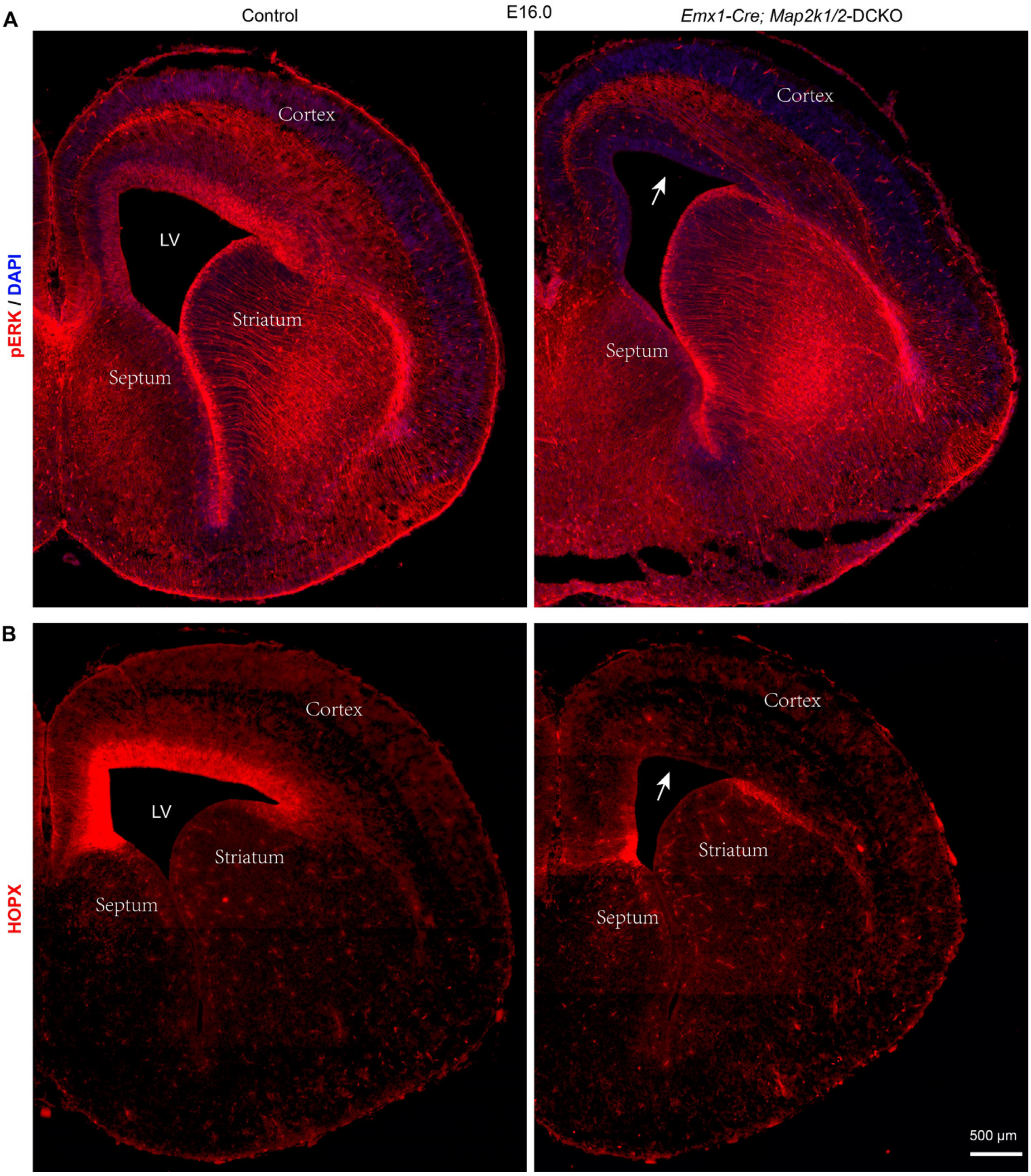


**Fig. S3. HOPX expression in cortical RGs depends largely on ERK signaling.** (***A***) pERK (phosphorylated ERK) immunostaining in E16.0 control mouse cortical RGs shows a medial-low to lateral-high gradient, reflecting higher YAP/TAZ activity medially, which represses both PKA and ERK signaling. pERK immunoreactivity is abolished in cortical RGs of *Emx1-Cre; Map2k1/2-dcko* mice at E16.0 (arrow). (***B***) In *Emx1-Cre; Map2k1/2-dcko* mice at E16.0, HOPX expression is specifically lost in the vast majority of cortical RGs (arrow) compared to controls. However, HOPX expression is preserved in the ventral pallium, where *Emx1-Cre* is not active in RGs, and in the lowermost medial cortical RGs, where BMP signaling may also promote its expression. Thus, HOPX expression levels in cortical RGs serve as a reliable readout of ERK signaling activity.

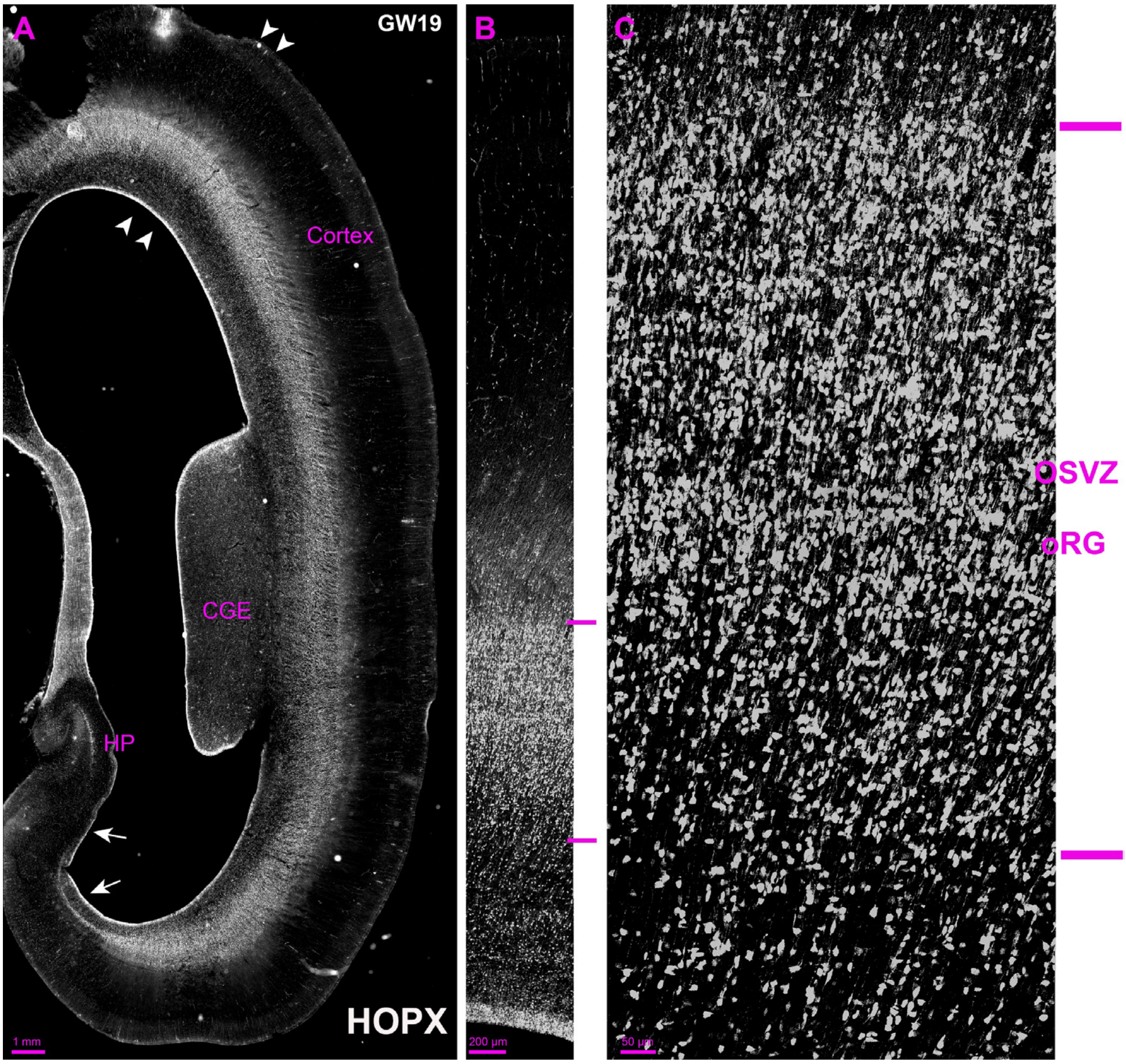


**Fig. S4. Human cortical oRGs express very higher HOPX at GW19. (*A*)** HOPX immunostaining of human GW19 telencephalic coronal sections at the posterior caudal ganglionic eminence (CGE) level. HOPX$^+$ oRGs were absent from the medial cortex (arrows), likely because elevated YAP/TAZ signaling in medial RGs promotes their conversion to E-tRGs and rapid differentiation into ependymal cells. HP, hippocampal region. (***B***) Higher-magnification view of the region indicated in (***A***) (arrowheads), further illustrating the HOPX expression pattern. HOPX$^+$ cells were distributed throughout the cortical region. (***C***) Higher-magnification view of the OSVZ in (***B***), highlighting strong HOPX expression in oRGs with larger soma.

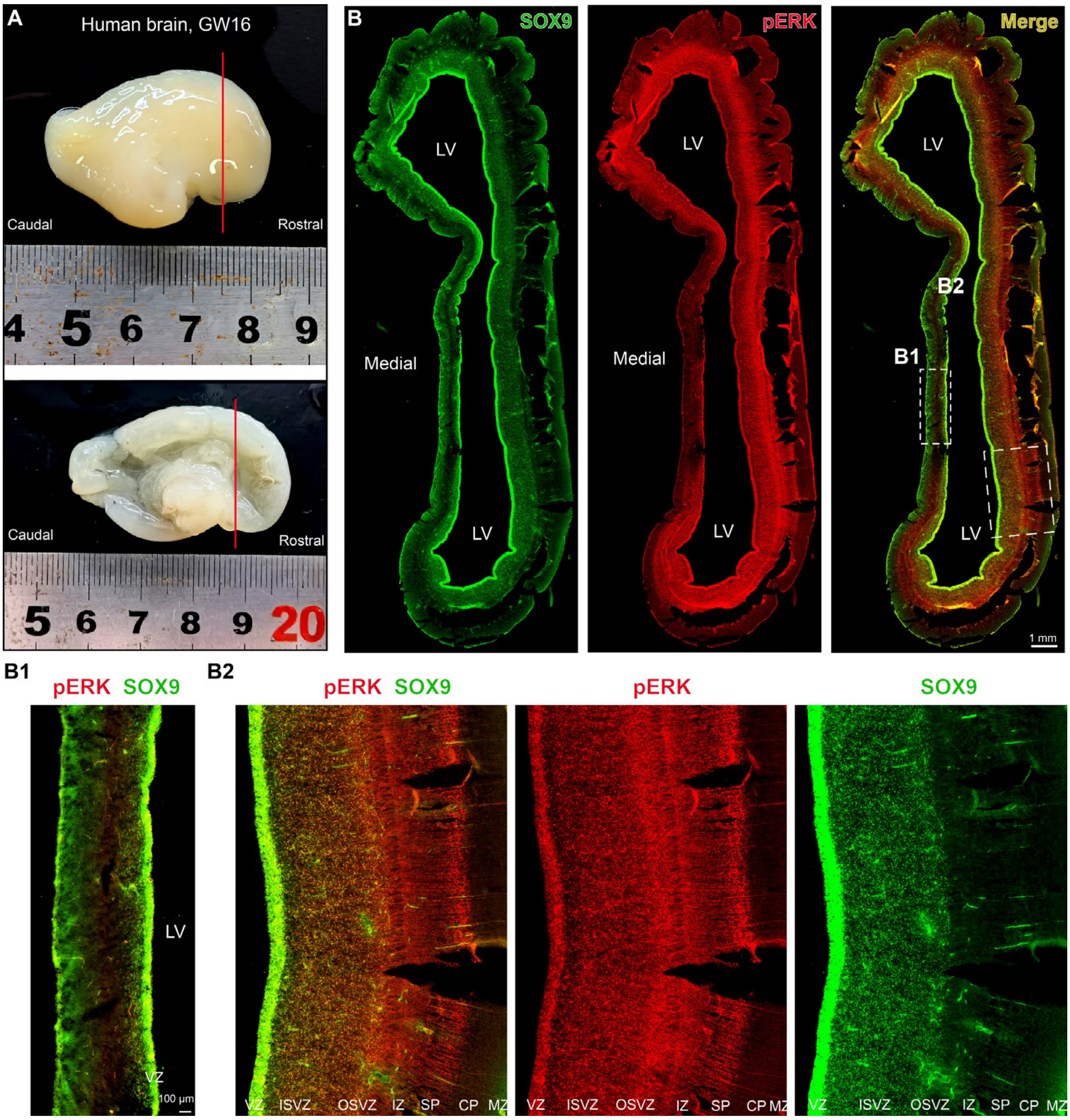


**Fig. S5. In the human GW16 medial cortex, oRGs are absent, whereas the dorsolateral and ventral cortices are enriched with oRGs and IPCs. (*A*)** Lateral and medial views of the GW16 human hemisphere. Red lines indicate the positions of the sections analyzed for immunofluorescence staining. **(*B*)** Double immunostaining for pERK and SOX9 on human cortical coronal sections at GW16. **(*B1*)** Higher-magnification view of the GW16 medial cortex, highlighting the absence of SOX9$^{+}$ IPCs and pERK$^{+}$/SOX9$^{+}$ oRGs in this region. **(*B2*)** Higher-magnification view of the GW16 lateral cortex, highlighting abundant SOX9$^{+}$ IPCs and pERK$^{+}$/SOX9$^{+}$ oRGs in the OSVZ. VZ, ventricular zone; ISVZ, inner subventricular zone; OSVZ, outer subventricular zone; IZ, intermediate zone; SP, subplate; CP, cortical plate; MZ, marginal zone.

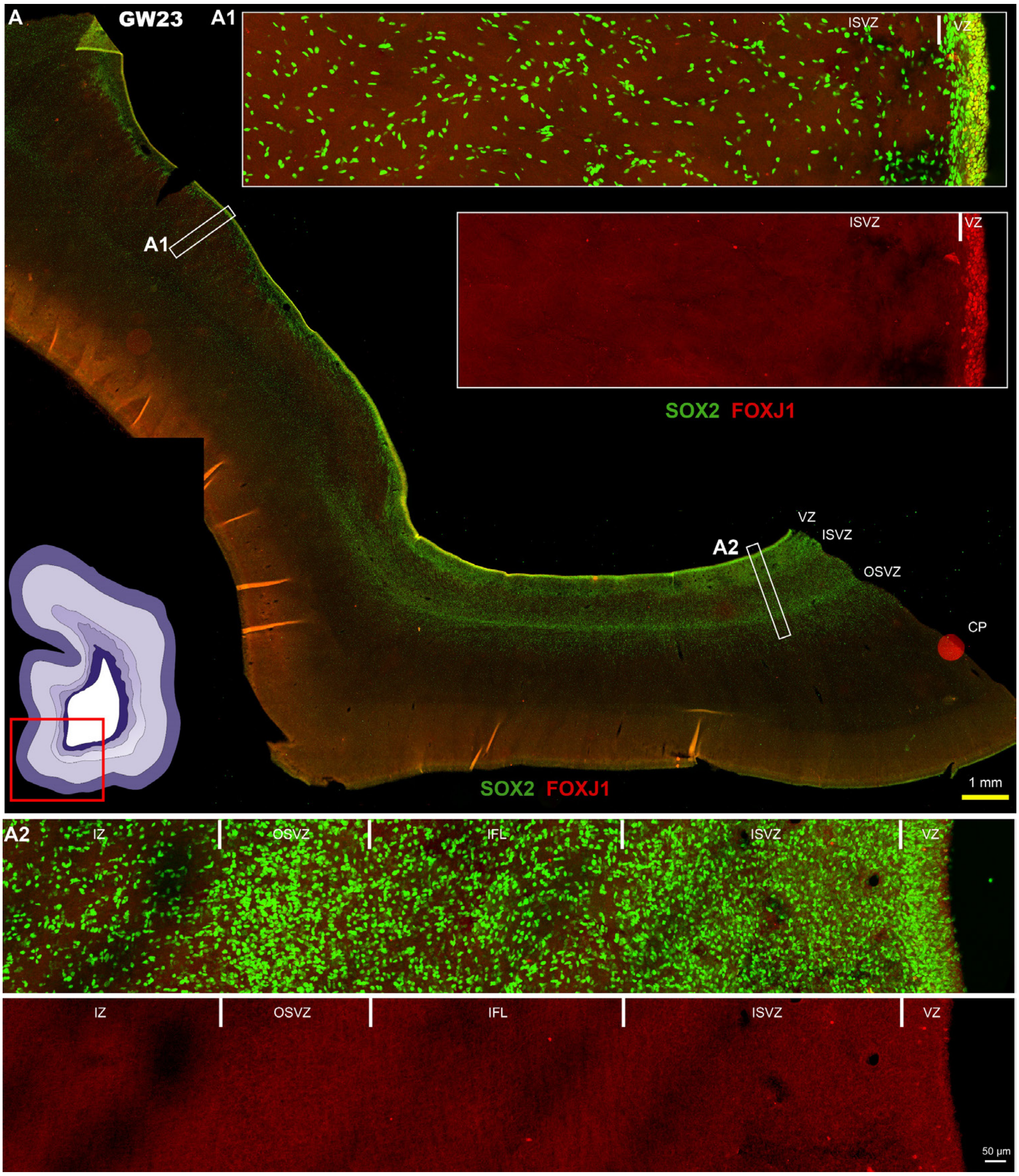


**Fig. S6. Representative immunofluorescence image of a human GW23 caudal cortex section, showing that oRGs are absent from the medial cortex but abundant in the ventrolateral cortices, where IPCs are also enriched. (A)** GW23 human cortical section, double-immunostained for SOX2 and FOXJ1. This image was previously published as Fig. S8 in our earlier study, where FOXJ1 staining alone was shown (12). The schematic shows the fragmented cortical section and its anatomical context. **(*A1*)** Higher-magnification view of the GW23 medial cortex, showing absent SOX2$^+$ IPCs but numerous FOXJ1$^+$/SOX2$^+$ E-tRGs and/or immature ependymal cells in the VZ. **(*A2*)** Higher-magnification view of the GW23 ventrolateral cortex, showing abundant SOX2$^+$ IPCs in the ISVZ and IFL, as well as abundant SOX2$^+$/FOXJ1$^-$ oRGs in the OSVZ and T-tRGs in the VZ. IFL, inner fibrous layer.

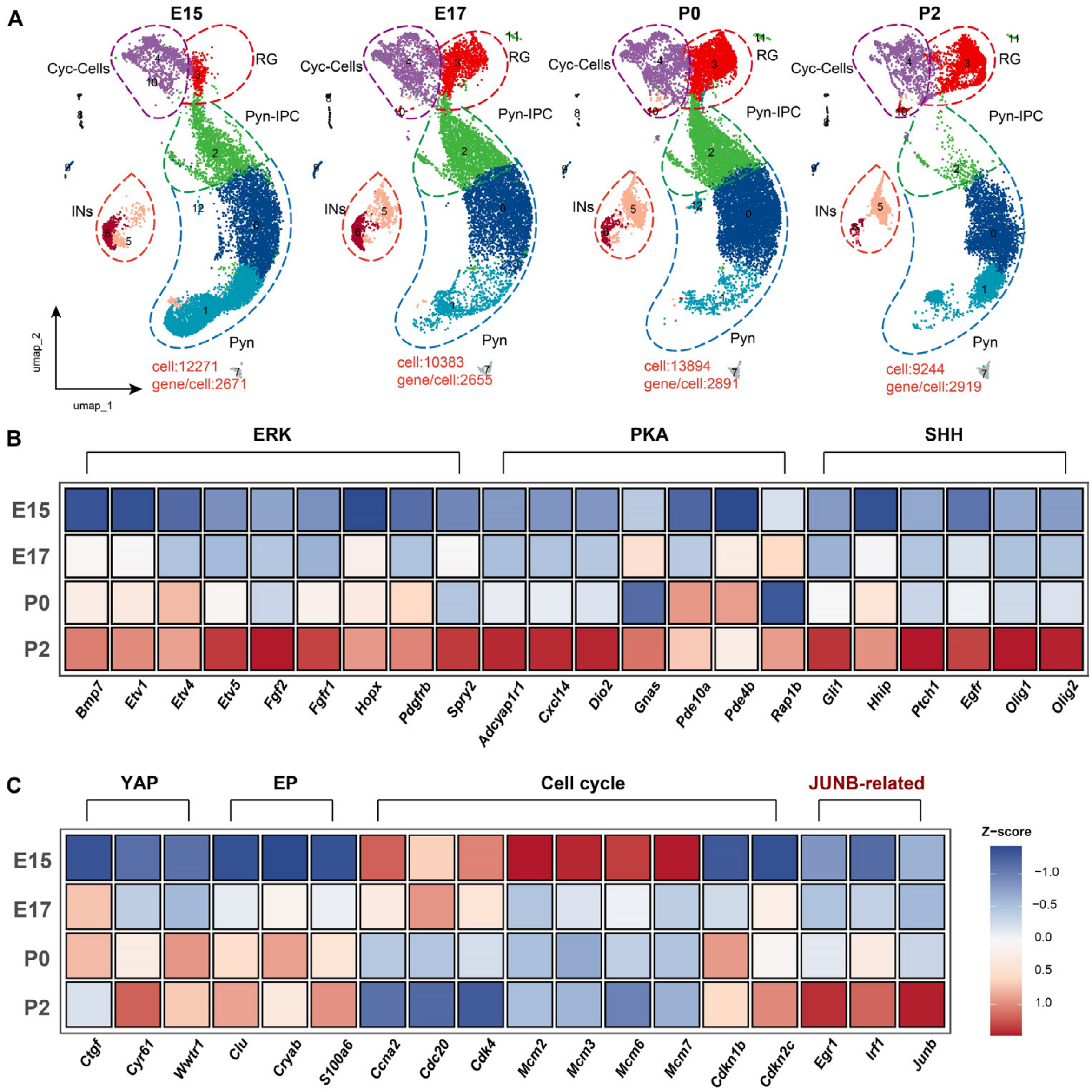


**Fig. S7. *Junb* is absent from mouse cortical RGs before E17.5, appears at P0, and reaches peak expression by P2, coinciding with the generation of ependymal cells and olfactory bulb interneurons (OBINs). (A)** Integrated high-quality scRNA-seq datasets from mouse cortices at E15.0, E17.0, P0, and P2, each containing abundant cortical RGs. The E15.0, E17.0, and P2 datasets were generated in our previous studies, while the P0 dataset is newly acquired in this work (see Methods). **(*B, C*)** From E15.0 to P2, cortical RGs (cluster 3 in ***A***) show progressive upregulation of ERK, PKA, SHH-SMO, and YAP/TAZ signaling, alongside early ependymal (EP) markers, reflecting cumulative pathway crosstalk. Concomitantly, cell cycle speed declines, faster at E15.0 than at P2. *Junb* is undetectable before E17.5, emerges at P0, and becomes strongly enriched by P2, coinciding with RG differentiation into ependymal cells and OBINs. *Egr1* and *Irf1*, the putative regulators of *Junb*, exhibited expression patterns largely consistent with that of *Junb.* Thus, *Junb* is not expressed in neurogenic mouse cortical RGs.

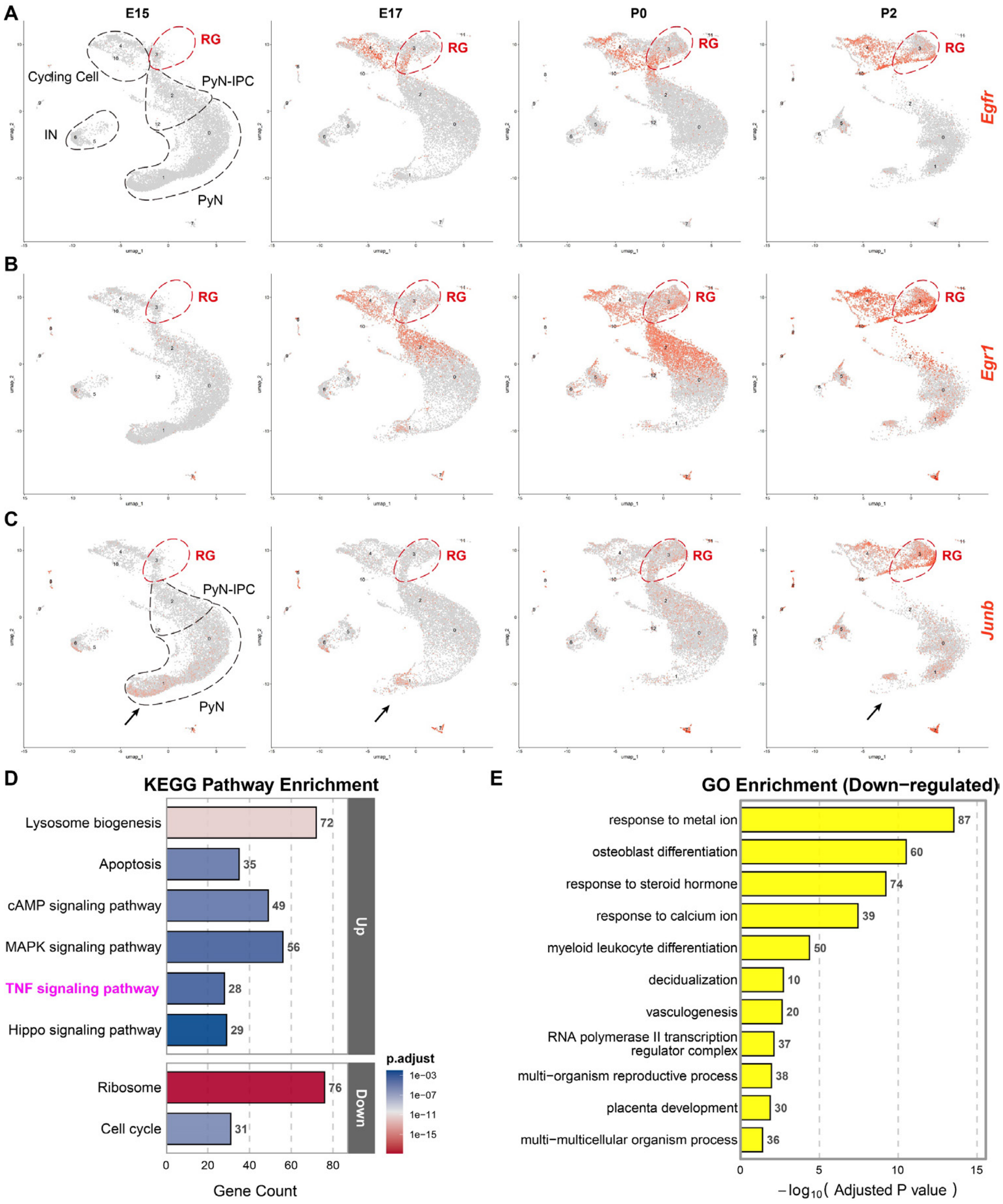


**Fig. S8. *Junb* expressed in later mouse cortical RGs might play a role in regulating key components of the TNF signaling pathway. (*A-C*)** Feature plots showing *Egfr*, *Egr1*, and *Junb* expression in mouse cortices at E15.0, E17.0, P0, and P2. All three genes exhibited progressively increasing expression levels in cortical RGs and IPCs during development, with low expression during the neurogenic period. Note *Junb* expression in a subset of mouse cortical PyNs **(**arrows in ***C*****)**. **(*D*)** KEGG analysis of genes upregulated in P2 versus E15.0 mouse cortical RGs revealed enrichment of the TNF pathway, which includes *Junb*. **(*E*)** GO analysis of *Junb*-associated genes upregulated in P2 versus E15.0 mouse cortical RGs revealed enrichment in transcriptional regulation, immune differentiation, and responses to hormonal and ionic stimuli, among others.

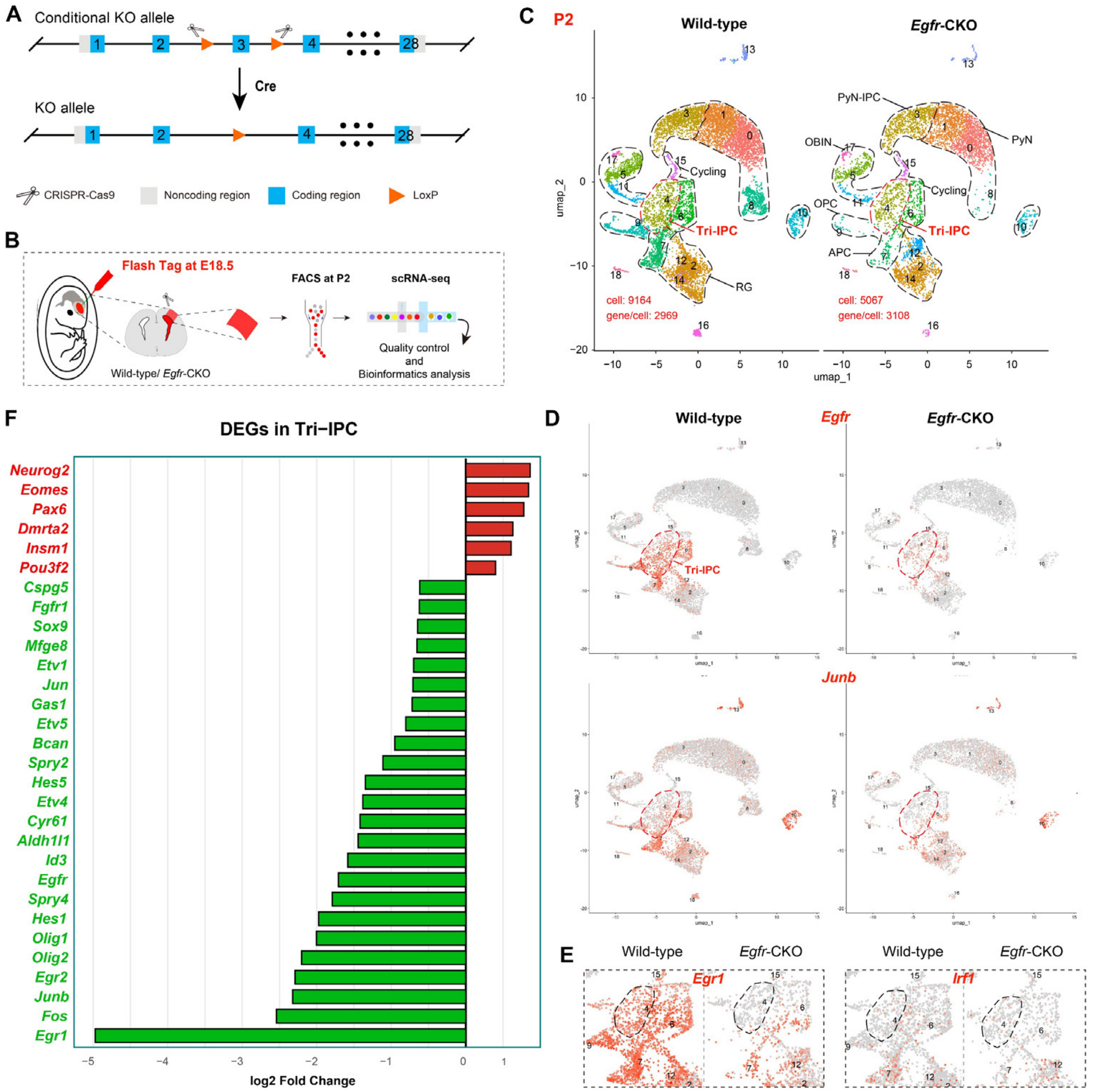


**Fig. S9. *Junb* expression in postnatal cortical Tri-IPCs is largely dependent on *Egfr*.** **(*A*)** Schematic of the *Egfr* floxed allele. Exon 3, which contains 184 bp of coding sequence, is flanked by loxP sites. Cre-mediated excision disrupts EGFR protein function (6). **(*B*)** scRNA-Seq of FACS-sorted cortical cells from P2 mouse littermates. FlashTag was administered at E18.5, and cortical cells were harvested at P2. **(*C*)** UMAP projection of scRNA-Seq data from P2 control and *hGFAP-Cre; Egfr-cko* mouse cortices, generated by re-analysis of our previously published datasets (6). **(*D, E*)** Feature plots of *Egfr, Junb, Egr1*, and *Irf1* in P2 control and *hGFAP-Cre; Egfr-cko* mouse cortical scRNA-Seq data. *Junb* and *Egr1* were highly expressed in Tri-IPCs, and both were markedly reduced in *hGFAP-Cre; Egfr-cko*, indicating that *Junb* and *Egr1* expression in postnatal cortical IPCs is largely *Egfr*-dependent. **(*F*)** Horizontal bar plot of $\log_2$ fold changes for DEGs in Tri-IPCs, showing upregulation of PyN-IPC marker genes and downregulation of EGFR-ERK signaling components and Tri-IPC markers. These findings suggest that EGFR promotes cortical gliogenesis while suppressing cortical neurogenesis (PyN-IPC genesis).

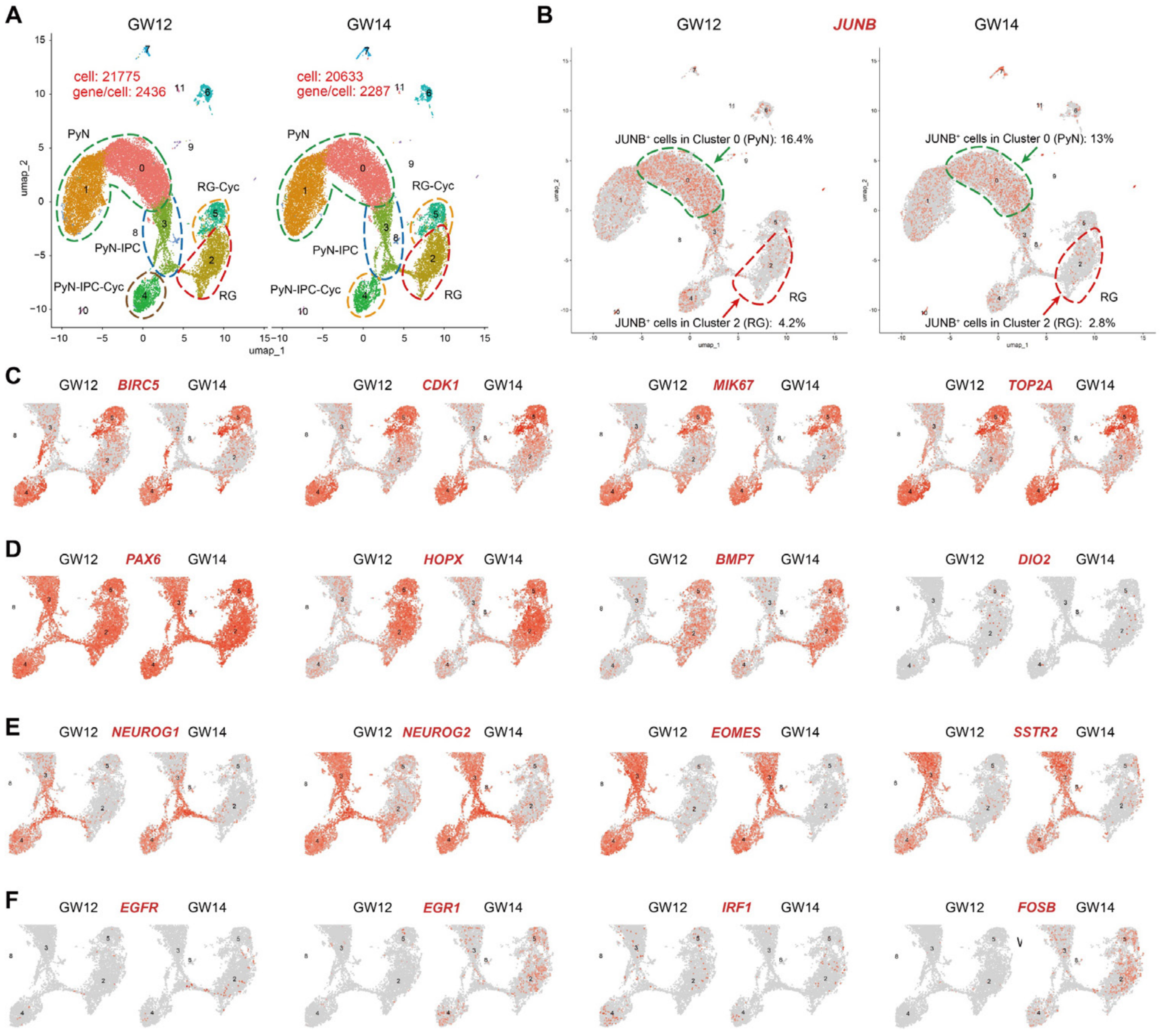


**Fig. S10. *JUNB* is barely detectable in human GW12 and GW14 cortical fRGs, which are exclusively undergoing cortical neurogenesis. (*A*)** UMAP projection of scRNA-Seq data from GW12 and GW14 human cortices, generated by re-analyzing previously published datasets (15). **(*B*)** Feature plots of *JUNB* expression in GW12 and GW14 human cortical scRNA-Seq data. *JUNB* was barely detectable in fRGs (4.2% at GW12; 2.8% at GW14) but was expressed in a subset of PyNs in cluster 0 (16.4% at GW12; 13% at GW14). This expression pattern is largely consistent with that observed in mouse. **(*C*)** Feature plots showing expression of proliferation marker genes (*BIRC5, CDK1, MKI67*, and *TOP2A*) in GW12 and GW14 human cortical scRNA-Seq data. **(*D*)** Expression patterns of *PAX6, HOPX, BMP7*, and *DIO2* in GW12 and GW14 human cortical fRGs. **(*E*)** Expression patterns of PyN-IPC marker genes (*NEUROG1, NEUROG2, EOMES*, and *SSTR*) in GW12 and GW14 human cortical scRNA-Seq datasets. **(*F*)** Expression patterns of *EGFR, EGR1, IRF1*, and *FOSB* in GW12 and GW14 human cortical fRGs. Overall, all four genes showed low expression at the neurogenic stages (GW12 and GW14), with substantially higher levels at later stages (GW17, GW18, and GW22–26) (See **Fig. S11-S14**).

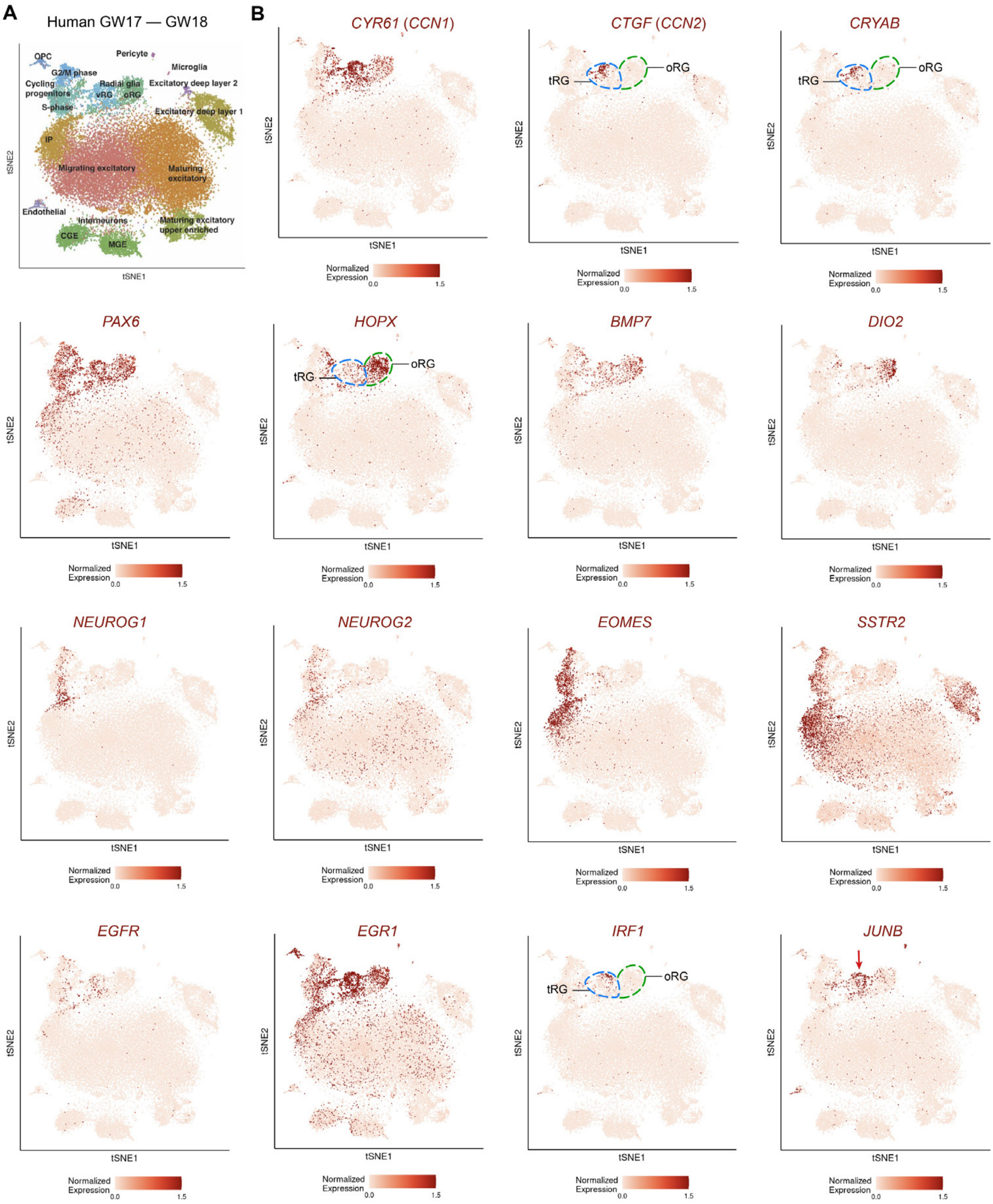


**Fig. S11. *JUNB* is expressed in GW17–GW18 human cortical tRGs but is rarely detected in oRGs. (*A*)** t-distributed stochastic neighbor embedding (tSNE) plot of GW17 and GW18 human cortical scRNA-seq data, with cells colored by cluster and annotated by major cell types according to the original study (16). Note that the cells originally designated as vRGs are now recognized as tRGs. At GW17 and GW18, human cortical RGs are simply categorized into two groups: oRGs and tRGs. **(*B*)** tSNE projections of oRG, tRG, and PyN-IPC marker genes show that *JUNB* is predominantly expressed in tRGs (arrow), with little to no expression in cortical oRGs at human GW17 and GW18.

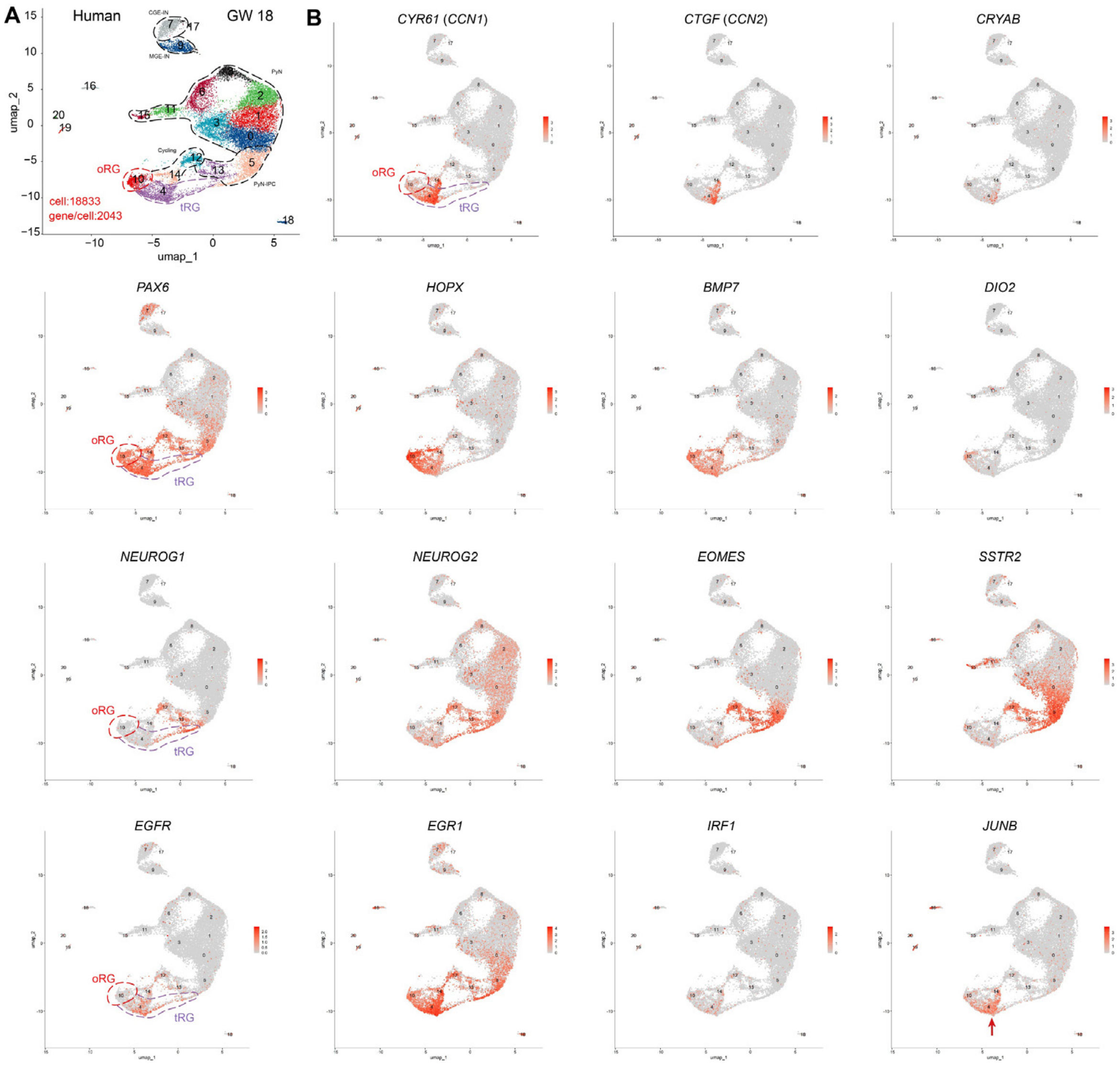


**Fig. S12. *JUNB* is selectively expressed in GW18 human cortical tRGs, with minimal expression in oRGs. (*A*)** UMAP of GW18 human cortical scRNA-seq data (published dataset) (17), with cells colored by cluster and labeled according to our reanalysis. At GW18, human cortical RGs fall into two simple categories: oRGs and tRGs. **(*B*)** *JUNB* expression in the GW18 human cortex is predominantly restricted to tRGs (arrow), with little to in oRGs, based on feature plots of oRG, tRG, and PyN-IPC markers.

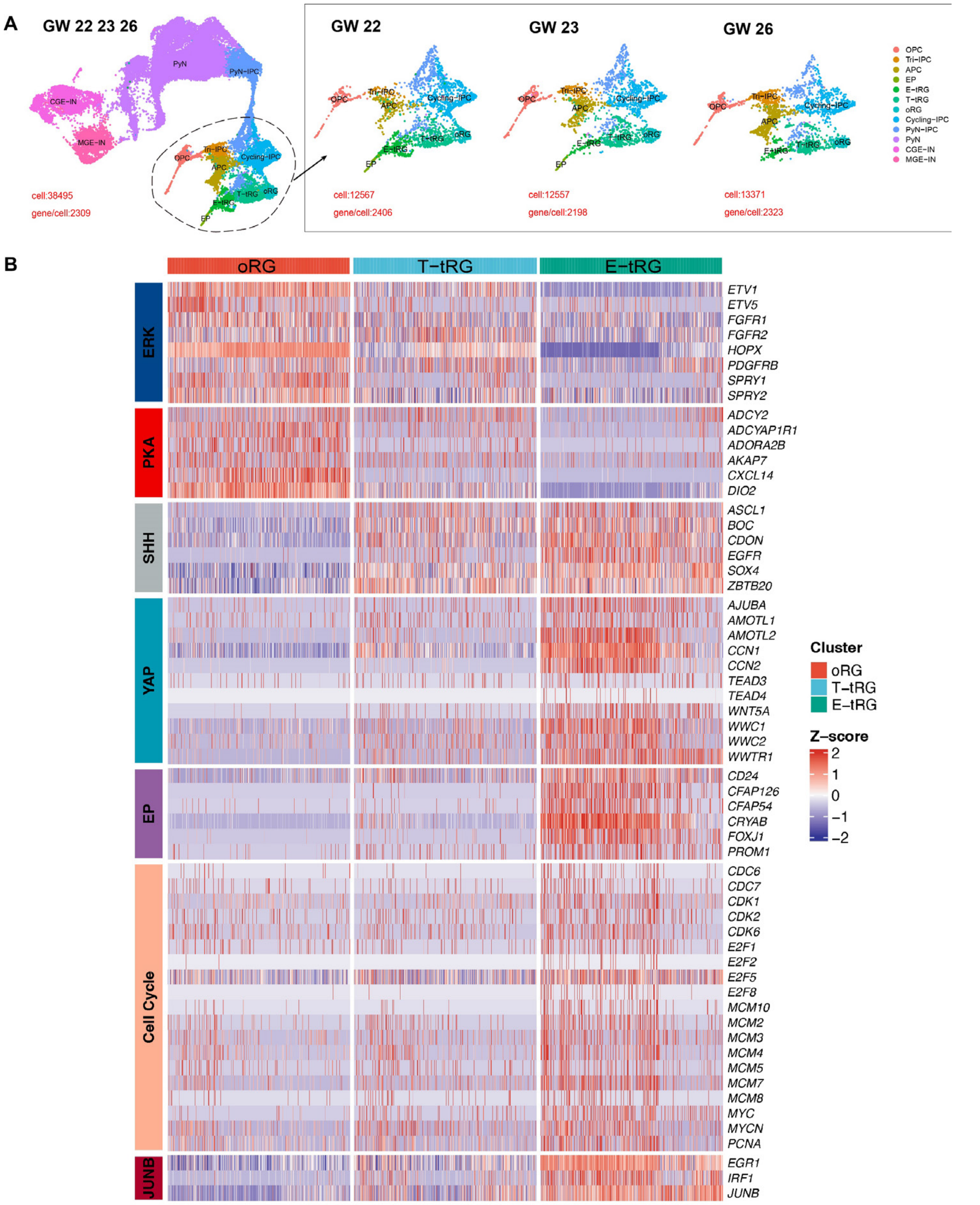


**Fig. S13. *JUNB* is highly expressed in E-tRGs, moderately in T-tRGs, and lowly in oRGs of the human cortex at GW22–GW26. (*A*)** UMAP visualization of scRNA-seq data from GW22, GW23, and GW26 human cortices (17). **(*B*)** Heatmap profiling reveals that oRGs exhibit heightened ERK/PKA activities. E-tRGs and T-tRGs show increasing SHH-SMO; however, in E-tRGs, high YAP/TAZ and FOXJ1 promote multiciliogenesis while suppressing SHH-SMO. Cell cycle genes are upregulated in T-tRGs and E-tRGs. Notably, in E-tRGs, the cell cycle program is repurposed for centriole amplification rather than mitotic division (19). JUNB is low in oRGs—reinforcing that JUNB is largely uninvolved in human cortical neurogenesis.

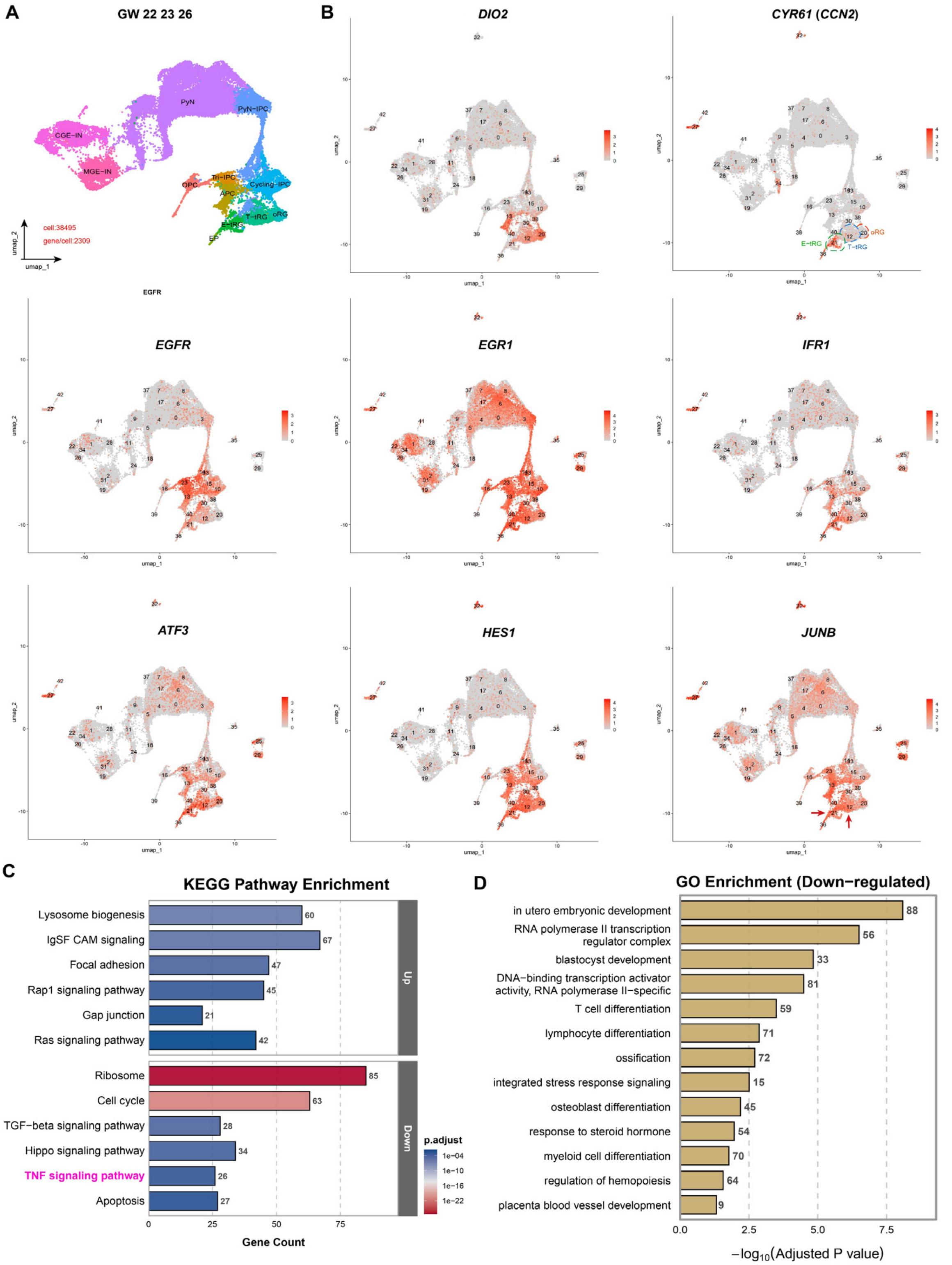


**Fig. S14. *JUNB* is expressed more in human cortical tRGs than oRGs and may regulate key components of the TNF pathway. (*A, B*)** Feature plots showing JUNB is enriched in E-tRGs and T-tRGs (arrows) over oRGs. **(*C*)** KEGG pathway analysis of genes upregulated in E-tRGs versus oRGs revealed enrichment of the TNF pathway, which includes *JUNB*. **(*D*)** GO analysis of *JUNB*-associated genes upregulated in E-tRGs versus oRGs revealed enrichment in transcriptional regulation, immune differentiation, and responses to hormonal and ionic stimuli, among others.

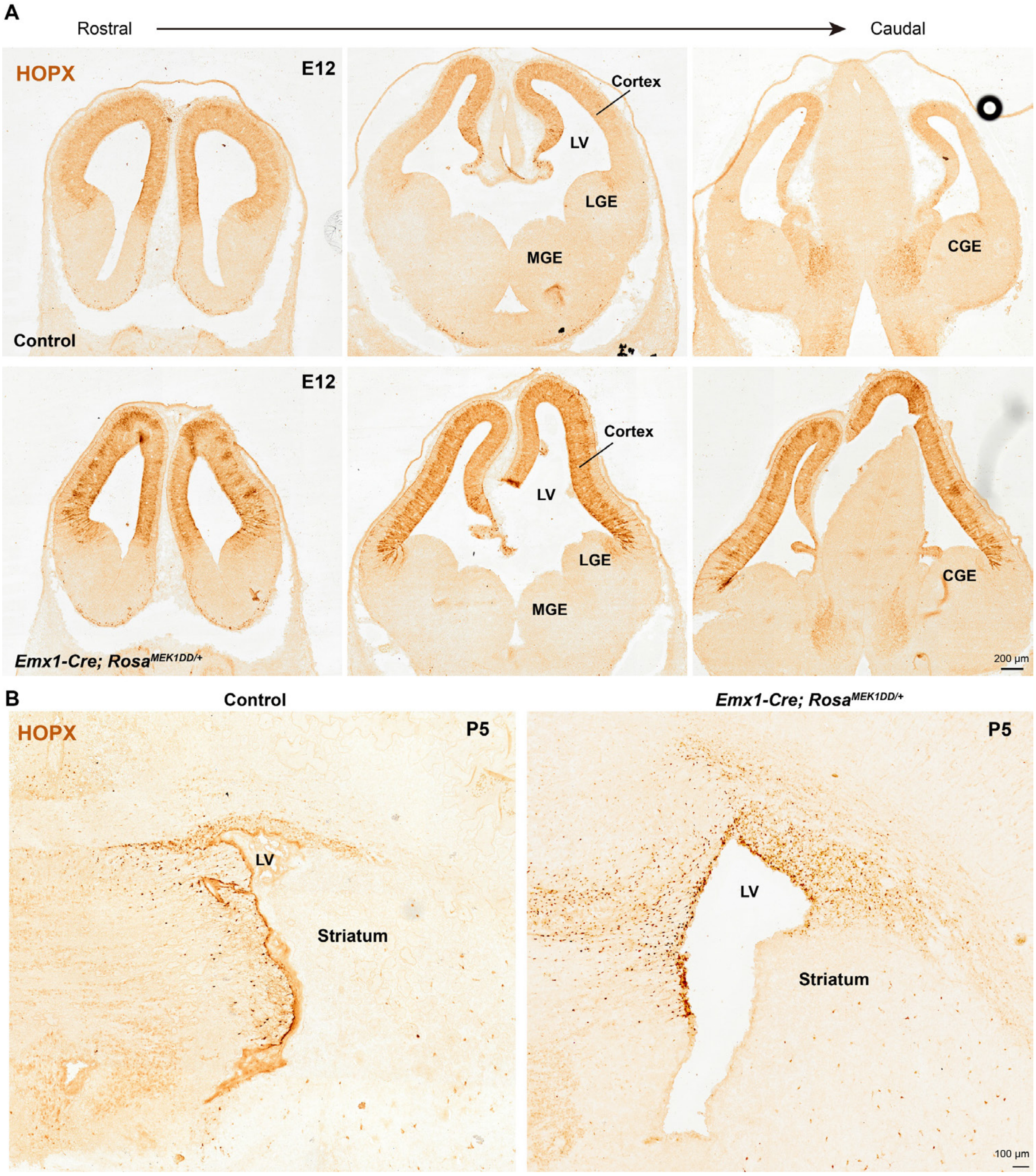


**Fig. S15. ERK signaling promotes HOPX expression in mouse cortical RGs. (*A*)** HOPX showed a rostral-high-to-caudal-low gradient in control cortical RGs at E12.0, mirroring the ERK activity gradient. *Emx1-Cre; Rosa26$^{MEK1DD/+}$* RGs exhibited elevated HOPX expression along the entire rostrocaudal axis at E12.0. **(*B*)** In *Emx1-Cre; Rosa26$^{MEK1DD/+}$* mice at P5, HOPX expression was elevated in cortical RGs-neural stem cells within the VZ, which corresponds to the *Emx1-Cre* expression domain.

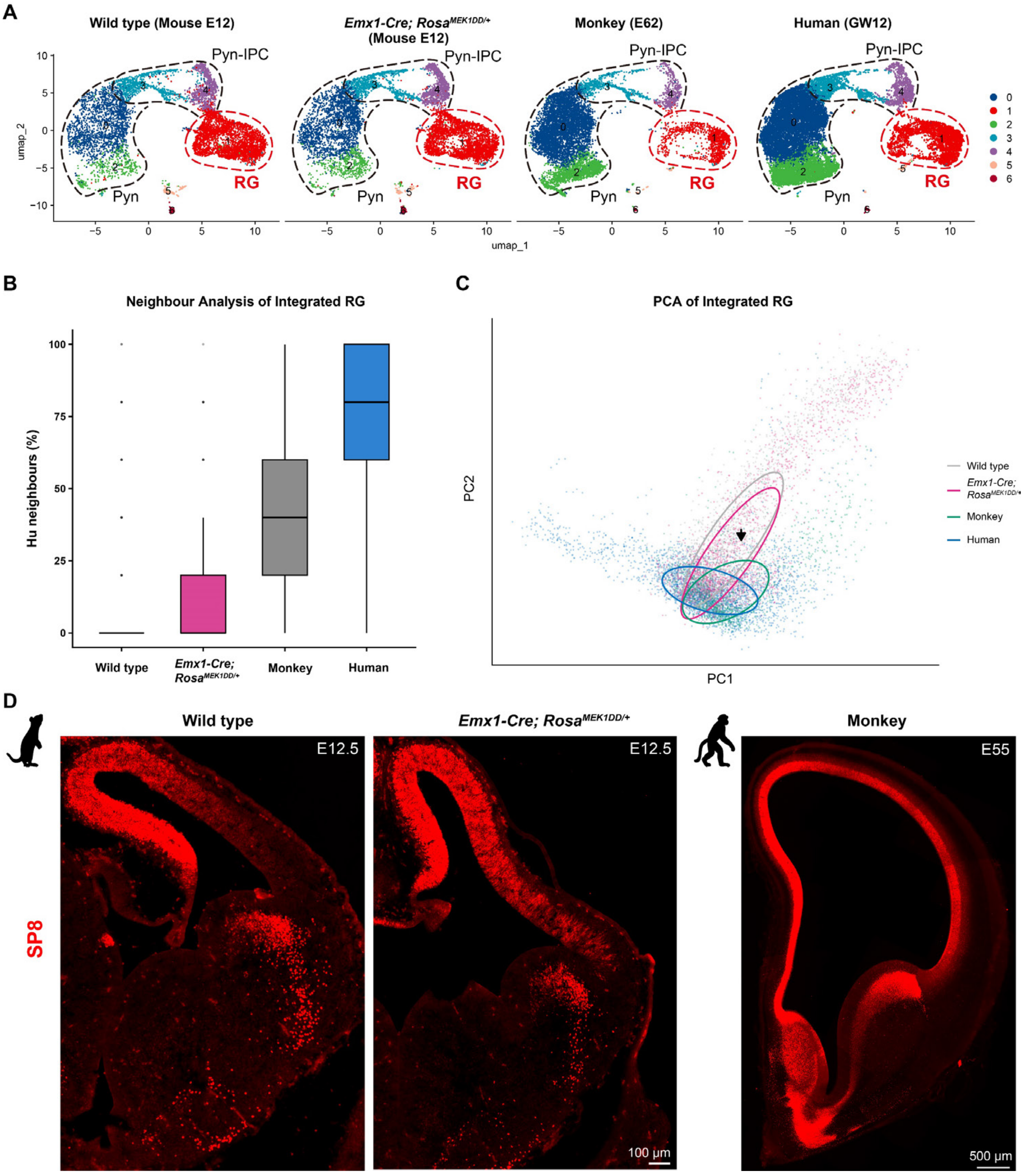


**Fig. S16. *MEK1DD* overexpression partially shifts the transcriptional landscape of early mouse cortical RGs toward that of primate cortical RGs. (*A*)** Integrated scRNA-seq analysis of mouse (E12.0, control and *Emx1-Cre; Rosa*$^{26MEK1DD/+}$), macaque (E62/E64) (18), and human (GW12) cortical datasets (15). **(*B, C*)** Neighbor-joining and principal component analyses suggested that *MEK1DD* overexpression partially shifts the mouse early cortical RG transcriptome toward a primate-like state. **(*D*)** SP8 immunofluorescence showed a graded expression pattern: strongest and most widespread in E55 macaque RGs, moderate in *Emx1-Cre; Rosa26*$^{MEK1DD/+}$ mouse RGs, and weakest in E12.0 control mouse RGs, supporting elevated ERK activity in macaque cortical RGs. The SP8-stained image of E55 macaque cortex was previously published as Fig. S10k in our earlier study (7), showing SP8/COUP-TFII double staining.

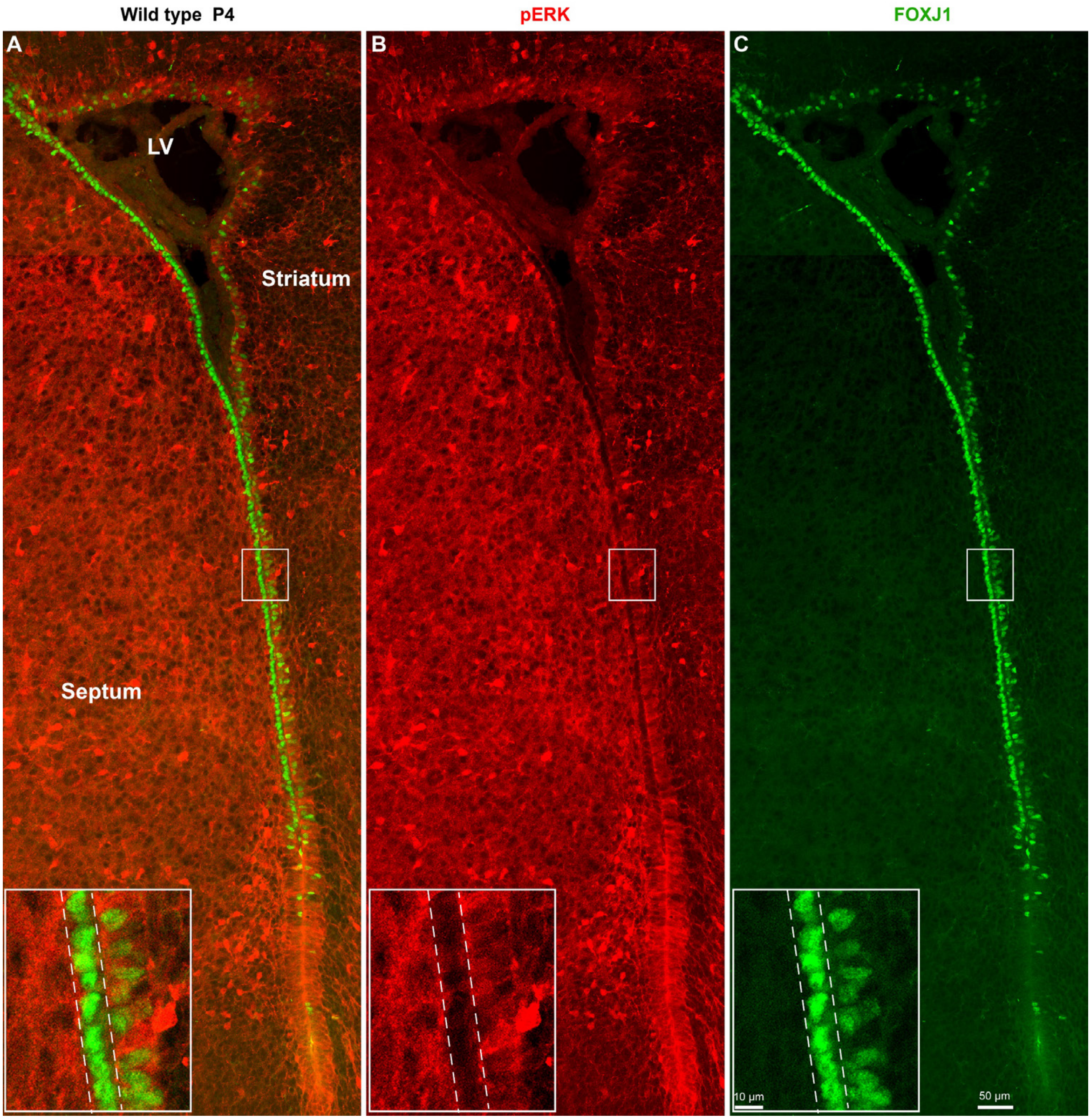


**Fig. S17. ERK expression and the ependymal differentiation program are mutually repressive in mouse telencephalic RGs/ependymal cells at P4. (*A–C*)** In P4 mouse brain sections double-immunostained for pERK and FOXJ1, pERK was consistently undetectable in the nuclei of FOXJ1-high cells, revealing a clear mutually repressive relationship between the two markers. This mutual inhibition between YAP/TAZ and ERK/PKA signaling accounts for the complete attenuation of ERK/PKA activity during YAP/TAZ-driven ependymal differentiation in immature ependymal cells.

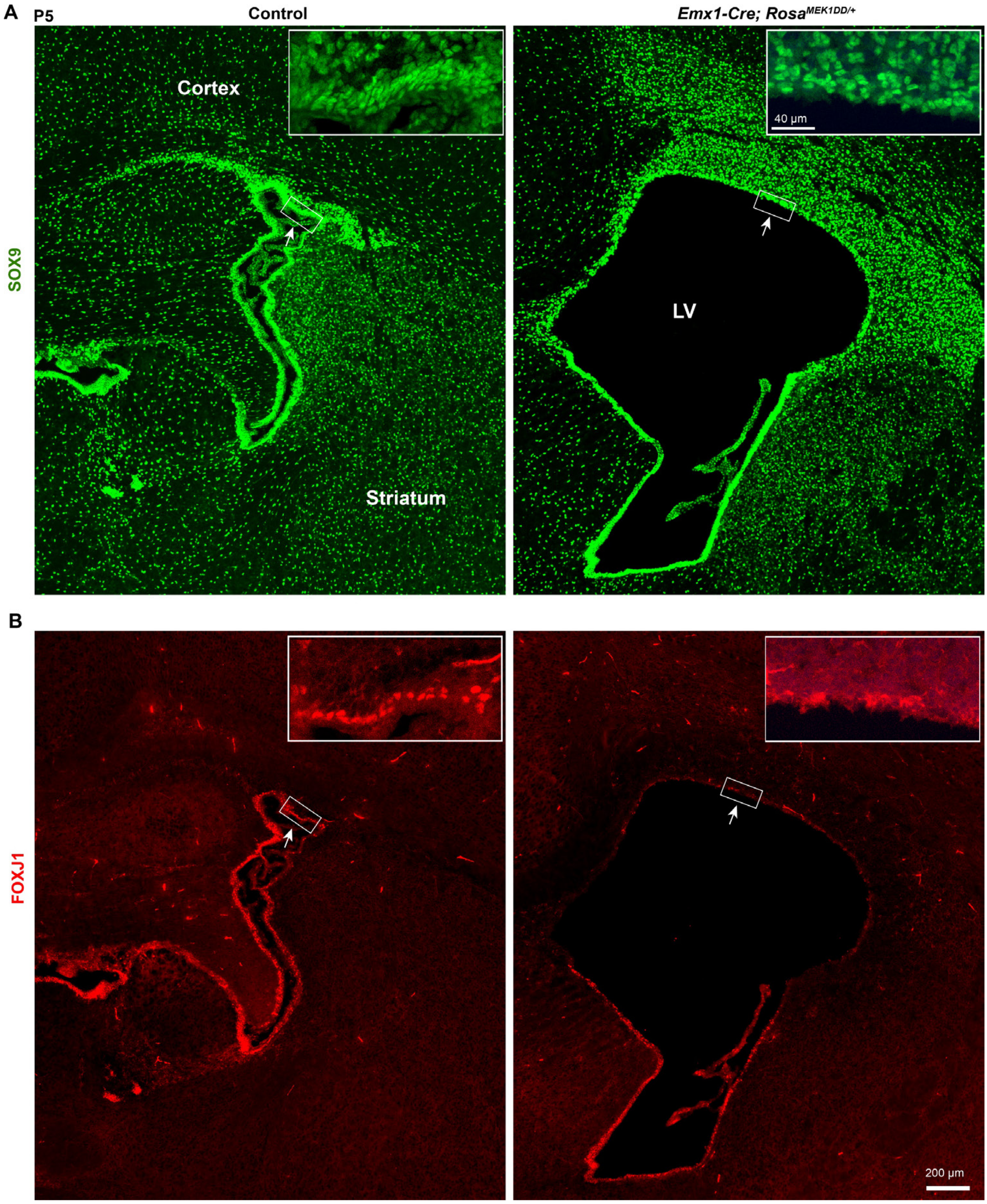


**Fig. S18. Sustained ERK overexpression blocks ependymal cell generation in mouse cortical RGs. (*A*)** SOX9 immunostaining revealed that sustained ERK activation in the *Emx1-Cre; Rosa26$^{MEK1DD/+}$* cortex led to extensive proliferation of SOX9$^{+}$ progenitors in the SVZ, accompanied by a marked loss of SOX9$^{+}$ ependymal cells in the VZ at P5 (see magnified VZ image in insets). **(*B*)** Sustained ERK activation in *Emx1-Cre; Rosa26$^{MEK1DD/+}$* cortical RGs completely blocked FOXJ1$^{+}$ ependymal cell generation within the *Emx1-Cre* expression domain at P5 (see magnified VZ image in insets).

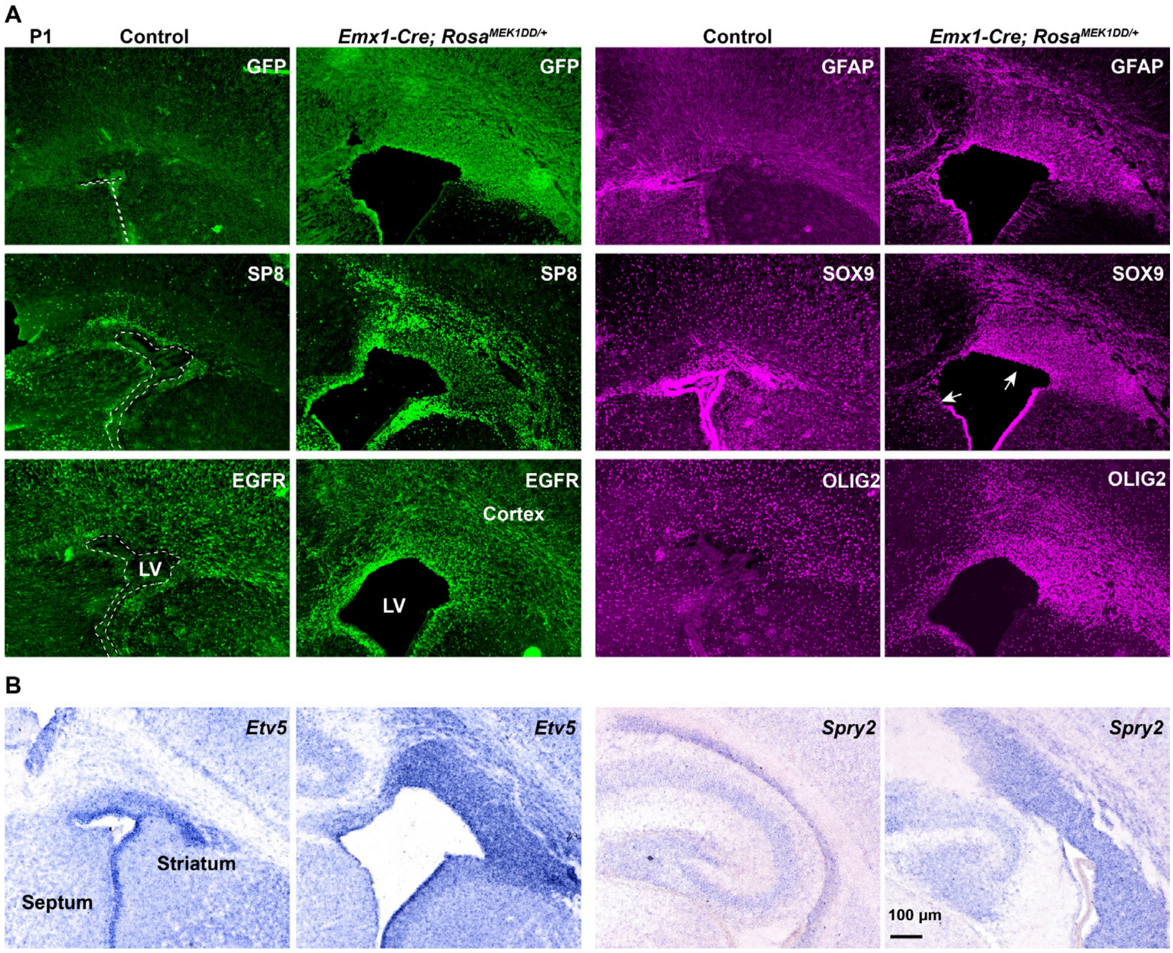


**Fig. S19. Sustained ERK overexpression in mouse cortex induces massive glial progenitor proliferation. (*A*)** P1 brain sections from control and *Emx1-Cre; Rosa26*$^{MEK1DD/+}$ mice were immunostained for MEK1DD-GFP, SP8 (OBIN-IPCs and OBINs), GFAP, SOX9, EGFR, and OLIG2. The mutant cortex showed extensive gliogenesis and increased OBIN production, suggesting that sustained ERK overexpression drives massive proliferation of Tri-IPCs and glial progenitors. Notably, ERK later upregulates EGFR expression, which may further potentiate EGFR-mediated ERK signaling. Arrows indicate loss of SOX9$^{+}$ immature ependymal cells in the VZ and disruption of the ependymal cell layer in *Emx1-Cre; Rosa26*$^{MEK1DD/+}$ mice. **(*B*)** In situ hybridization showed that *Etv5* and *Spry2* were markedly upregulated in cortical progenitors of P1 *Emx1-Cre; Rosa26*$^{MEK1DD/+}$ mice relative to controls, indicating that both genes are reliable transcriptional readouts of ERK signaling in the developing cortex.

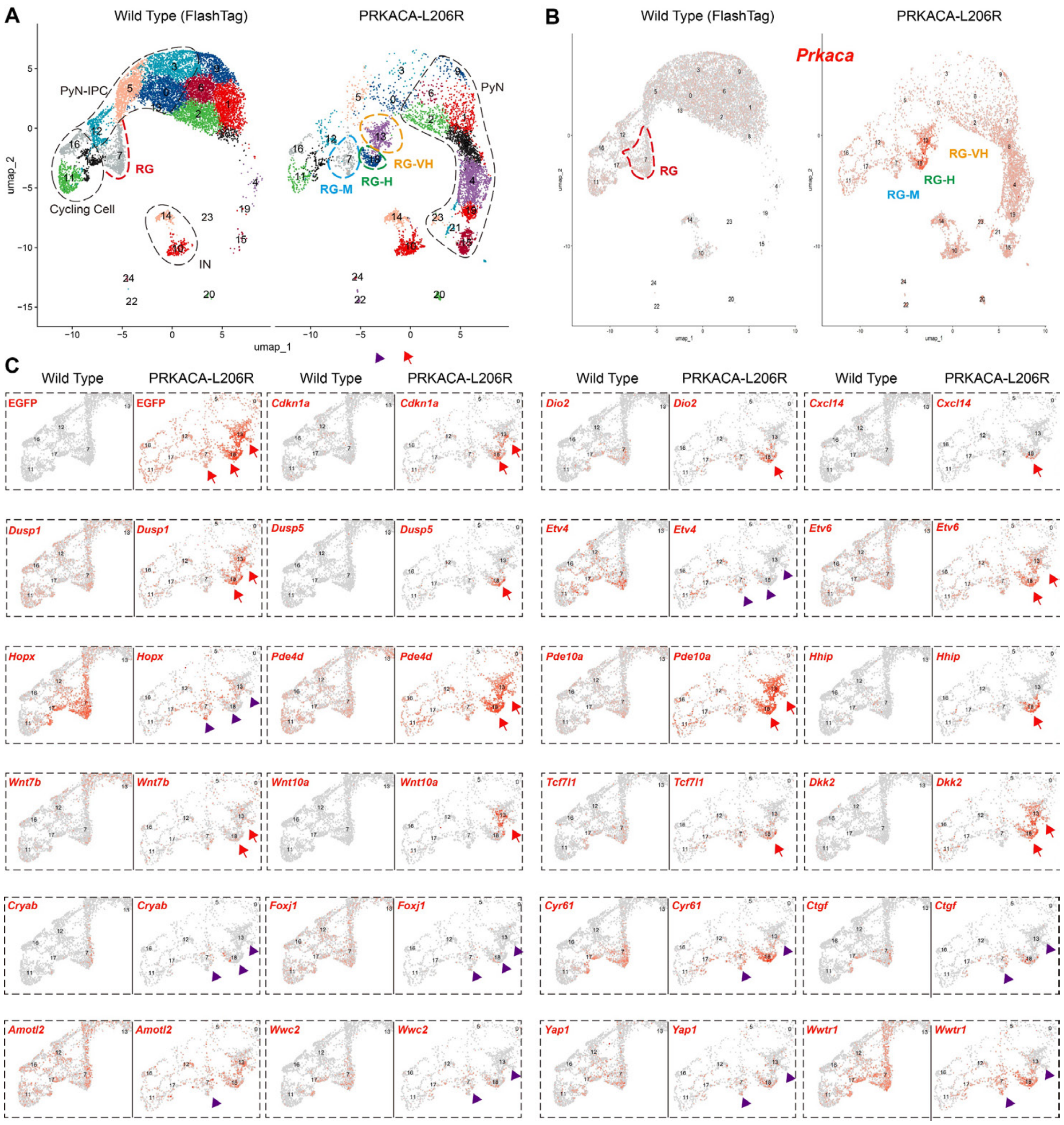


**Fig. S20. Gene expression profiles from E17.0 scRNA-seq data.** (***A-C***) UMAP of E17.0 cortical cells colored by cluster (same as Fig. 4B). *PRKACA-L206R-GFP*-labeled RGs were segregated into three clusters based on GFP and *Prkaca* intensity: RG-M (moderate, cluster 7), RG-H (high, cluster 18), and RG-VH (very high, cluster 13). Feature plots comparing gene expression in control RGs (cluster 7) and PRKACA-L206R-GFP$^{+}$ RG-M, RG-H, and RG-VH, respectively. *Cdkn1a* upregulation in RG-H and RG-VH slows the RG cell cycle. *Dio2* and *Cxcl14* upregulation in RG-H marks PKA signaling. Upregulation of *Dusp1/5* and *Etv6* and downregulation of *Etv4/Hopx* indicated strongly reduced ERK signaling. *Pde4d* and *Pde10a* upregulation indicates strong PKA signaling and represents negative feedback. *Hhip* upregulation in RG-H is likely driven by enhanced TGF-β signaling. Upregulation of *Wnt7b*, *Wnt10a*, *Tcf7l1*, and *Dkk2* likely reflects activation of both canonical and non-canonical Wnt signaling. *Cryab, Foxj1, Cyr61 (Ccn1), Ctgf (Ccn2), Wwc2, Yap1,* and *Wwtr1* were downregulated in RG-VH, indicating that very high PKA activity strongly inhibits YAP/TAZ signaling in cortical RGs. Arrows indicate gene upregulation; arrowheads indicate gene downregulation.

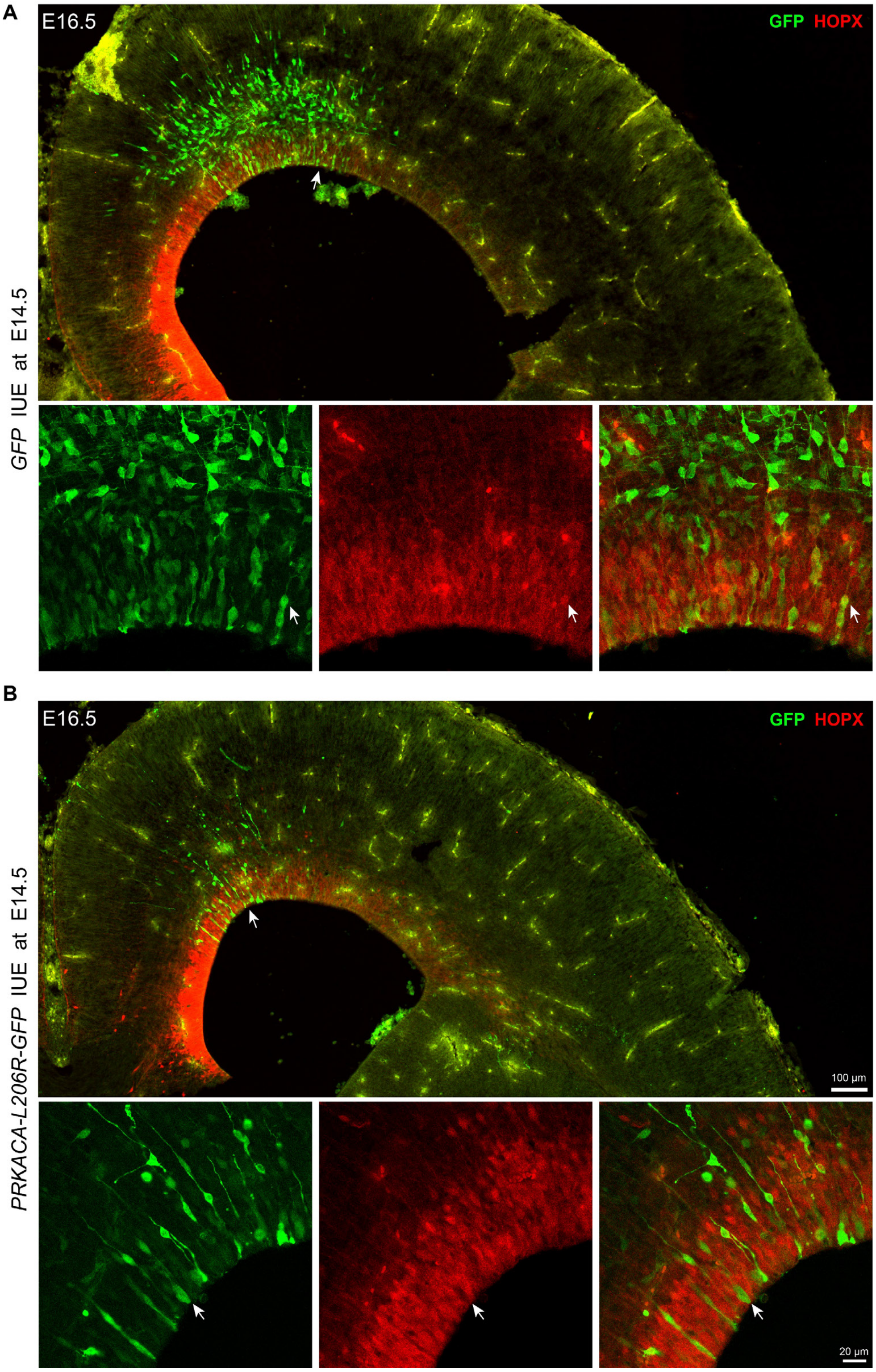


**Fig. S21. PKA overexpression suppresses HOPX expression in mouse cortical RGs.** (***A, B***) E14.5 IUE with *GFP* or *PRKACA-L206R-GFP*; analyzed at E16.5. GFP/HOPX co-immunostaining revealed that HOPX expression was nearly undetectable in PRKACA-L206R-GFP$^+$ cells (arrowhead), whereas GFP$^+$/HOPX$^+$ cells were readily observed in controls (arrows). Moreover, mutant RGs produced substantially fewer progeny than controls, consistent with PKA overexpression-mediated cell cycle slowing in cortical RGs.

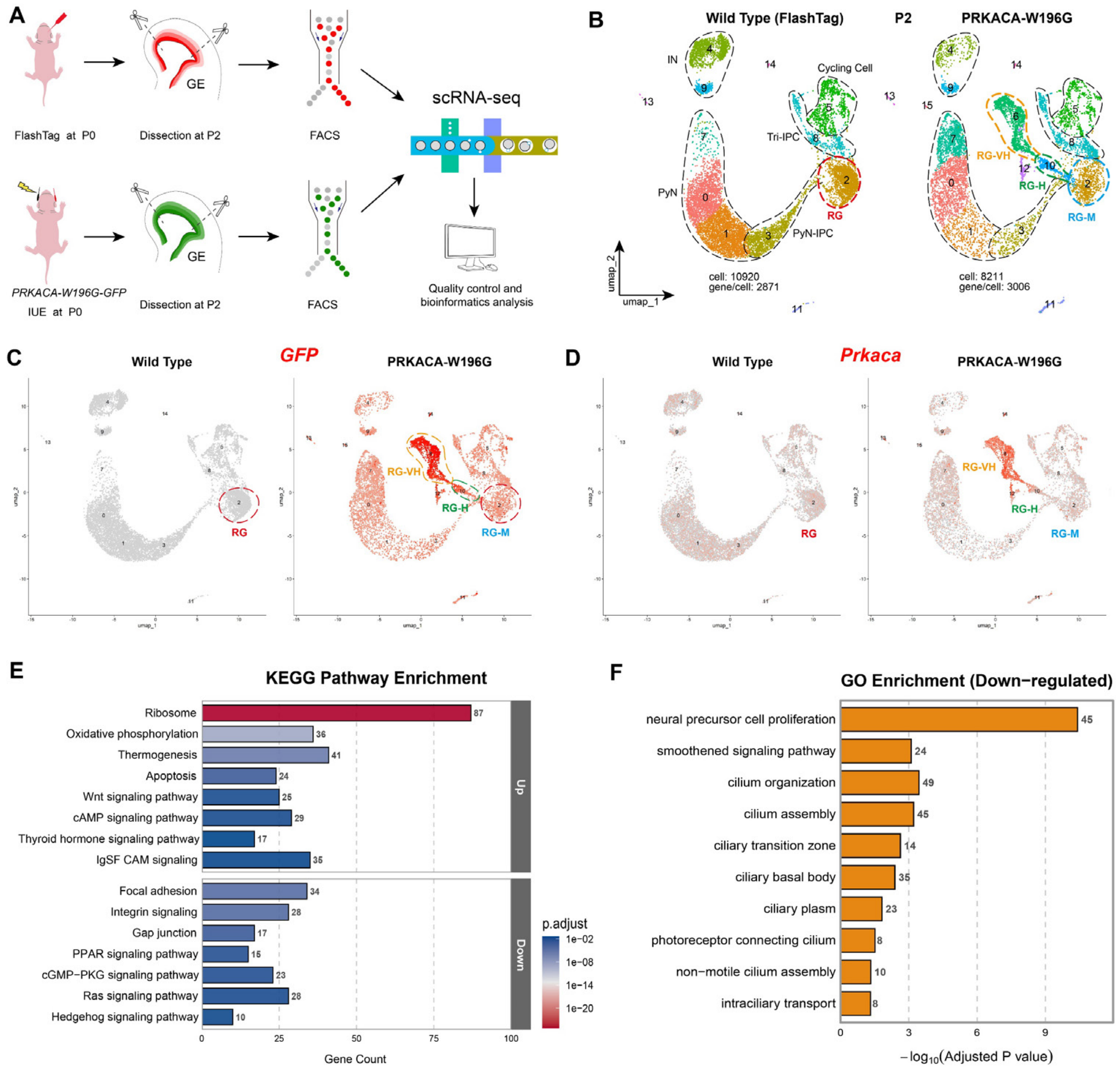


**Fig. S22. PKA overexpression in neonatal mouse cortical RGs suppresses SHH–SMO signaling and blocks ciliogenesis. (*A*-*D*)** scRNA-Seq of FACS-sorted cortical cells from P2 mouse littermates. Controls received FlashTag at P0; experimental embryos received *PRKACA-W196G-GFP* by IUE. Cells were collected 48 h later for sequencing. *PRKACA-W196G-GFP*-labeled RGs segregated into three clusters by GFP intensity: RG-M (moderate, cluster 2), RG-H (high, cluster 10), and RG-VH (very high, cluster 6). (***E***) KEGG analysis of DEGs between *PRKACA-W196G-GFP*-labeled RG-H and control RGs (WT cluster 2) revealed their involvement in multiple signaling pathways. Upregulated genes were enriched in cAMP signaling, apoptosis, and canonical and non-canonical WNT signaling, whereas downregulated genes were enriched in Integrin, Gap junction, cGMP-PKG, Ras, and SHH-SMO signaling. (***F***) Downregulated genes in P2 *PRKACA-W196G-GFP*-labeled RG-H versus controls (WT cluster 2) were enriched for ciliogenesis GO terms, implicating PKA overexpression in ciliogenesis suppression. Genes associated with neural precursor cell proliferation and SHH–SMO signaling were also downregulated.

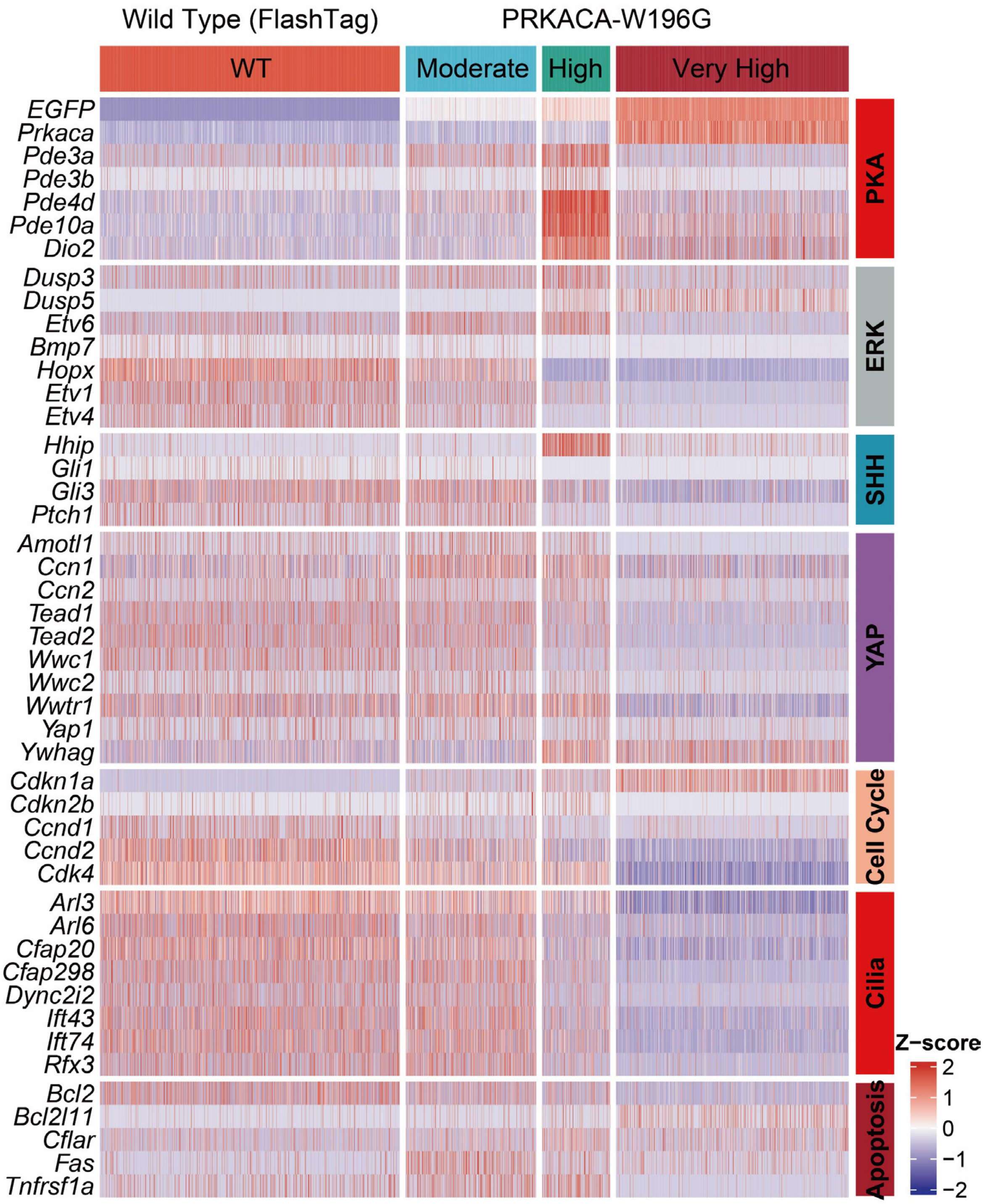


**Fig. S23. PKA overexpression elicits widespread functional effects in mouse cortical RGs.** Heatmap of DEGs in P2 cortical RGs labeled with *PRKACA-W196G* (RG-M, RG-H, and RG-VH) versus controls. PKA overexpression dose-dependently suppressed ERK, SHH-SMO, and YAP/TAZ signaling, slowed the cell cycle, blocked ciliogenesis, and induced apoptosis.

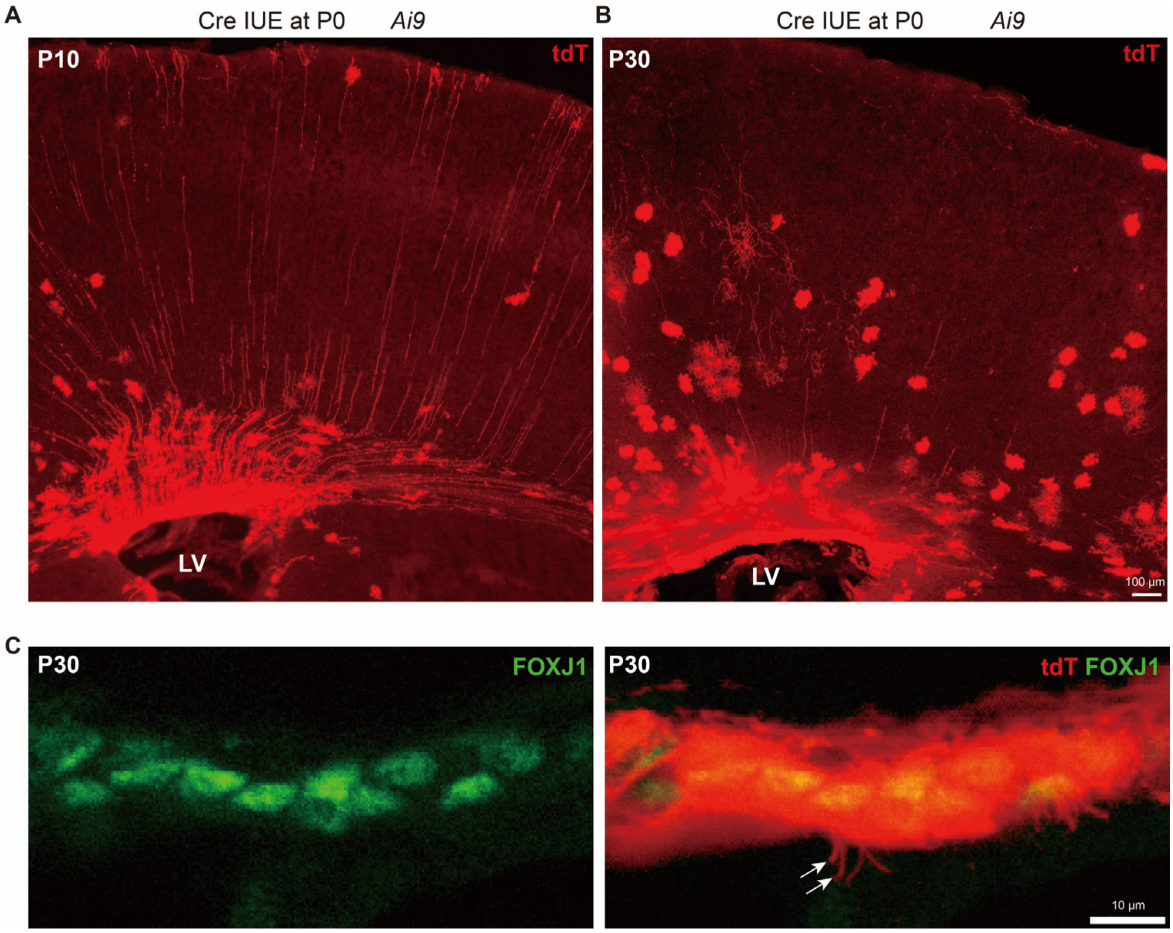


**Fig. S24. Electroporation of *Cre*-expressing plasmids at P0 efficiently labeled cortical RGs and their descendants in *Ai9* mice. (*A*)** *Cre*-expressing plasmids were electroporated into cortical RGs of P0 *Ai9* mice. tdT (tdTomato)-labeled cortical RGs were traceable to P10, with a subset retaining full-span morphology. **(*B*)** At P30, numerous tdT-labeled cortical astrocytes and oligodendrocytes were identified based on their glial morphology. **(*C*)** High-magnification imaging at P30 identified FOXJ1$^+$/tdT$^+$ mature multiciliated ependymal cells in the cortical VZ.

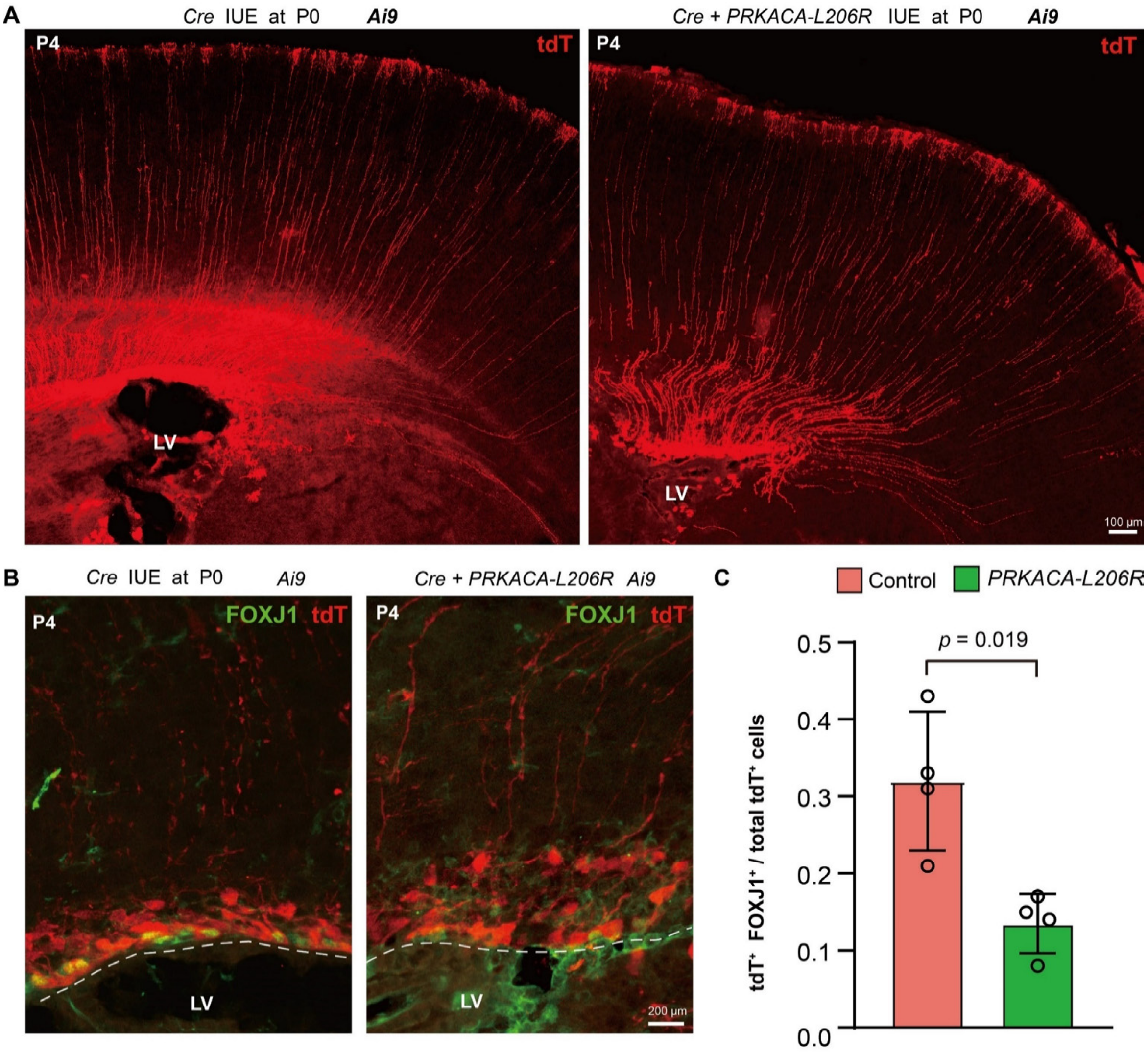


**Fig. S25. PKA overexpression in neonatal mouse cortical RGs blocks ependymal cell generation.** (***A–C***) P0 *Ai9* cortical RGs were electroporated with *Cre* (control) or *Cre* + *PRKACA-L206R-GFP* (experimental). Tissues were harvested at 4 days post-electroporation. In both groups, tdT-labeled cortical RGs maintained intact full-span morphology. However, tdT/FOXJ1 immunostaining revealed significantly fewer tdT$^{+}$/FOXJ1$^{+}$ cells in the VZ of PRKACA-L206R samples compared with controls. Moreover, the ependymal layer was disrupted and lacked an intact lining in experimental tissues.

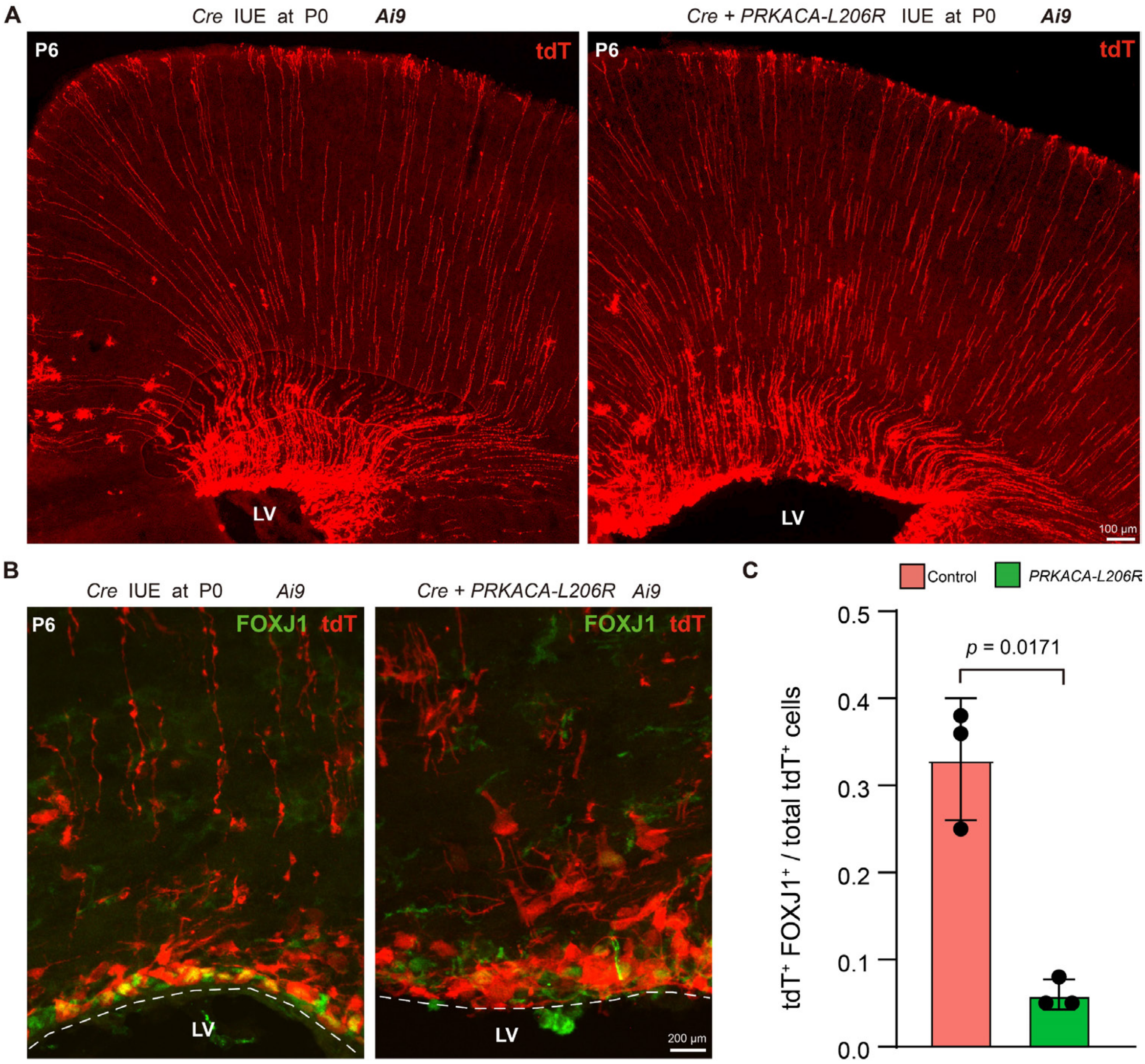


**Fig. S26. PKA overexpression in neonatal mouse cortical RGs prevents ependymal cell generation.** (***A–C***) P0 *Ai9* cortical RGs were electroporated with *Cre* (control) or *Cre* + *PRKACA-L206R-GFP* (experimental), and harvested at 6 days. tdT-labeled RGs retained intact full-span morphology in both groups. However, tdT/FOXJ1 staining revealed fewer $tdT^{+}/FOXJ1^{+}$ VZ cells in mutants versus controls.
